# Dopamine neurons of the VTA encode active conspecific interaction and promote social learning through social reward prediction error

**DOI:** 10.1101/2020.05.27.118851

**Authors:** Clément Prévost-Solié, Benoit Girard, Beatrice Righetti, Malika Tapparel, Camilla Bellone

## Abstract

Social interactions motivate behavior in many species, facilitating learning, foraging and cooperative behavior. However, how the brain encodes the reinforcing properties of social interactions remains elusive. Here using in vivo recording in freely moving mice, we show that Dopamine (DA) neurons of the Ventral Tegmental Area (VTA) increase their activity during active interactions with unfamiliar conspecific. Using a social instrumental task, we then show that VTA DA neuron activity signals social reward prediction error and drives social reinforcement learning. Thereby, our findings propose that VTA DA neurons are a neural substrate for a social learning signal driving motivated behavior.

**One Sentence Summary:** DA neurons are a substrate for social reward learning through the Social Reward Prediction Error.

## Main Text

Dopamine neurons of the Ventral Tegmental Area (VTA DA neurons) are at the core of the reward system and critical for reward processing and exploration of novel stimuli (*1*) (*2*). Electrophysiological studies support the hypothesis that DA neurons signal Reward Prediction Error, the difference between the expected and the actual reward (*3*-*5*). This signal is needed to update reward expectations and facilitate learning (*6*) and impaired reward prediction error encoding may underlie the pathophysiology of several psychiatric disorders.

Especially, growing body of literature hypothesizes that deficits in reward prediction error in social context may explain social deficits in psychiatric disorders (*7*) but whether DA neurons support social prediction is still largely unknown.

Over the past decade, some evidences suggested the rewarding nature of social interaction (*8, 9*) (*10*) and neuroimaging studies and fiber photometry analysis have recently demonstrated that social stimuli recruit the reward system (*11, 12*). In this study we investigated which aspects of the interaction with a conspecific are rewarding and whether the activity of VTA DA neurons guides social reinforcement learning via social reward prediction error (SRPE).

Using *in-vivo* recordings in freely behaving mice (Suppl. Fig. 1A-D), we recorded the activity of DA neurons during direct interaction with an unfamiliar conspecific (here defined as social stimulus; Fig. 1A, B). Experimental DAT-Cre mice were first injected with AAV-DIO-ChR2 and implanted with a recording electrode in the VTA to identify and record DA neurons (Suppl. Fig. 1E-K) (*13*). Based on waveform and firing pattern of recorded neurons (Suppl. Fig. 1L-N), an unsupervised cluster analysis was performed and two spatially distinct groups were identified: non-putative DA and putative DA + photolabeled DA (Suppl. Fig. 1O-Q). Afterwards, we built a classification model reaching 99.6% of accuracy to predict cluster membership of the recorded neurons (Suppl. Fig. 1R). Only putative DA (pDA) neurons predicted by the model were used in the following analyzes.

**Figure 1.**
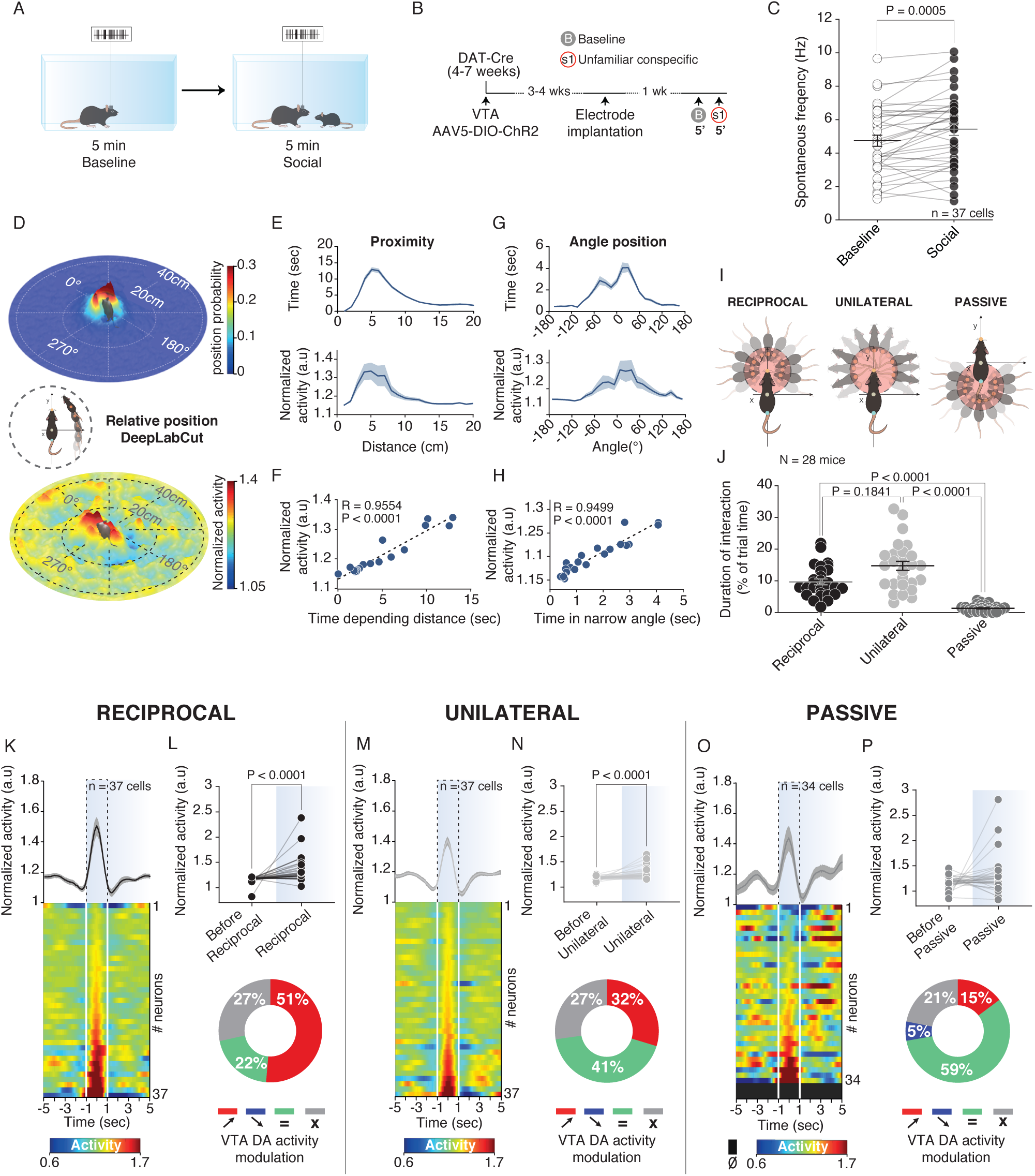
VTA DA neurons increase activity during free social interaction with high heterogeneity of response. **(A)** Schema of the free social interaction task. **(B)** Experimental time-course of the free social interaction. The DAT-Cre mice injected with an AAV-DIO-ChR2 and implanted with recordings electrodes in the VTA prior to perform the free social interaction task. **(C)** VTA DA spontaneous frequency at baseline and during social context. Paired t-test (t_(36)_ = -3.8275). (D) Middle: Schema for the calculation of the relative position of the social stimulus. Top: 3D heatmap of the relative position of the social stimulus. Down: 3D heatmap of the normalized VTA DA activity depending on the relative position of the social stimulus. **(E)** Top: Proximity; Time spent depending on the distance from the social stimulus (mean = 8.3230cm, SEM = 1.2633cm). Down: Normalized VTA DA activity depending on the distance from the social stimulus (mean = 7.906cm, SEM = 1.1079cm). **(F)** Normalized VTA DA activity depending on the time spent for each bin (1cm) of distance from the social stimulus. Linear regression with Pearson’s correlation coefficient (R = 0.9554, P < 0.0001). **(G)** Top: Angle position; Time spent depending on the angle made between the experimental mouse and the social stimulus (mean = 7.3205°, SEM = 32.6890°). Down: Normalized VTA DA activity depending on the angle made between the experimental mouse and the social stimulus (mean = -5.8675°, SEM = 22.4467°). **(H)** Normalized VTA DA activity depending on the time spent for each bin (10°) of angle between the experimental mouse and the social stimulus. Linear regression with Pearson’s correlation coefficient (R = 0.9499, P < 0.0001). **(I)** Schema of reciprocal (Left), unilateral (Middle) and passive (Right) interactions. **(J)** Time of interaction in reciprocal, unilateral or passive contacts during a same session. Friedman test (χ^2^_(28)_ = 45.50, P < 0.0001), followed by Dunn’s test. **(K)** Normalized VTA DA activity during reciprocal interaction. Top: PETH of the normalized VTA DA activity centered on the reciprocal interaction. Down: PETH’s heatmap of normalized VTA DA activity during reciprocal interaction for each neuron recorded. **(L)** Top: Comparison between the VTA DA activity before and during reciprocal interaction. Wilcoxon test (W = 619). Down: Proportion of VTA DA neurons increasing (red), decreasing (blue) or not changing their activity (green) during reciprocal interaction. The grey labelling indicates non-responding neurons independently the type of interaction (reciprocal, unilateral or passive). **(M)** Normalized VTA DA activity during unilateral interaction. Top: PETH of the normalized VTA DA activity centered on the unilateral. Down: PETH’s heatmap of normalized VTA DA activity during unilateral interaction for each neuron recorded. **(N)** Top: Comparison between the VTA DA activity before and during unilateral interaction. Wilcoxon test (W = 617). Down: Proportion of VTA DA neurons increasing, decreasing or not changing their activity during unilateral interaction. **(O)** Normalized VTA DA activity during passive interaction. Top: PETH of the normalized VTA DA activity centered on the passive interaction. Down: PETH’s heatmap of normalized VTA DA activity during passive interaction for each neuron recorded. **(P)** Top: Comparison between the VTA DA activity before and during passive interaction. Wilcoxon test (W = 125, P = 0.2932). Down: Proportion of VTA DA neurons increasing, decreasing or not changing their activity during passive interaction

During the 5 min of free interaction with a social stimulus, we observed an overall increase in the VTA DA spontaneous frequency and in the spikes within bursts (Fig. 1C; Suppl. Fig 2C, D).

Since accurate quantification of the behavior is necessary for understanding the neuronal correlates of social interaction, we used DeepLabCut (*14*) to analyze the relative position of the social stimulus toward the experimental mice (Fig.1D). We observed that the stimulus spends the majority of the time in the visual field and in the proximity of the experimental mouse (average proximity of 8.3 cm; Fig. 1E, G)). Furthermore, both the proximity and the angle position were strongly correlated with VTA DA activity (Fig. 1F, H), suggesting that VTA DA firing is associated to orientation and interaction toward the social stimulus.

To analyze the quality of conspecific interactions, we quantified active (reciprocal and unilateral) and passive contacts (Fig. 1I). Experimental mice spent more time engaging in reciprocal and unilateral compared to passive contacts (Fig. 1J). We observed that, although individual neuronal responses show high heterogeneity (Suppl. Fig. 2A, B), only active contacts induce an overall increase of VTA DA neuron firing (Fig. 1K-P) which is reflected in the VTA DA bursting activity (Suppl. Fig. 2E-P). The heterogeneity of neuronal responses suggests that VTA DA populations can be differentially recruited during interactions in social context. As a control, we considered behavior non-related to conspecific interaction and we observed that rearing did not increase VTA DA neuron activity (Suppl. Fig. 3A-C). Finally, we examined whether VTA DA neuron activity was sustained during the total duration of interaction bouts. Contrary to our prediction, the analysis revealed that stronger increase of VTA DA activity was correlated with shorter interactions. These experiments may suggest that longer bouts of interactions with a conspecific are not necessarily more rewarding or that DA neuron signal specific aspects of the interaction. Remarkably, we also observed that phasic increase was not sustained and the peak always occurred during the first half of each bouts of interaction for all the different contacts (Supp. Fig. 2Q-V). These experiments suggest that VTA DA neurons encode the initiation of the interaction more than the consumption.

**Figure 2.**
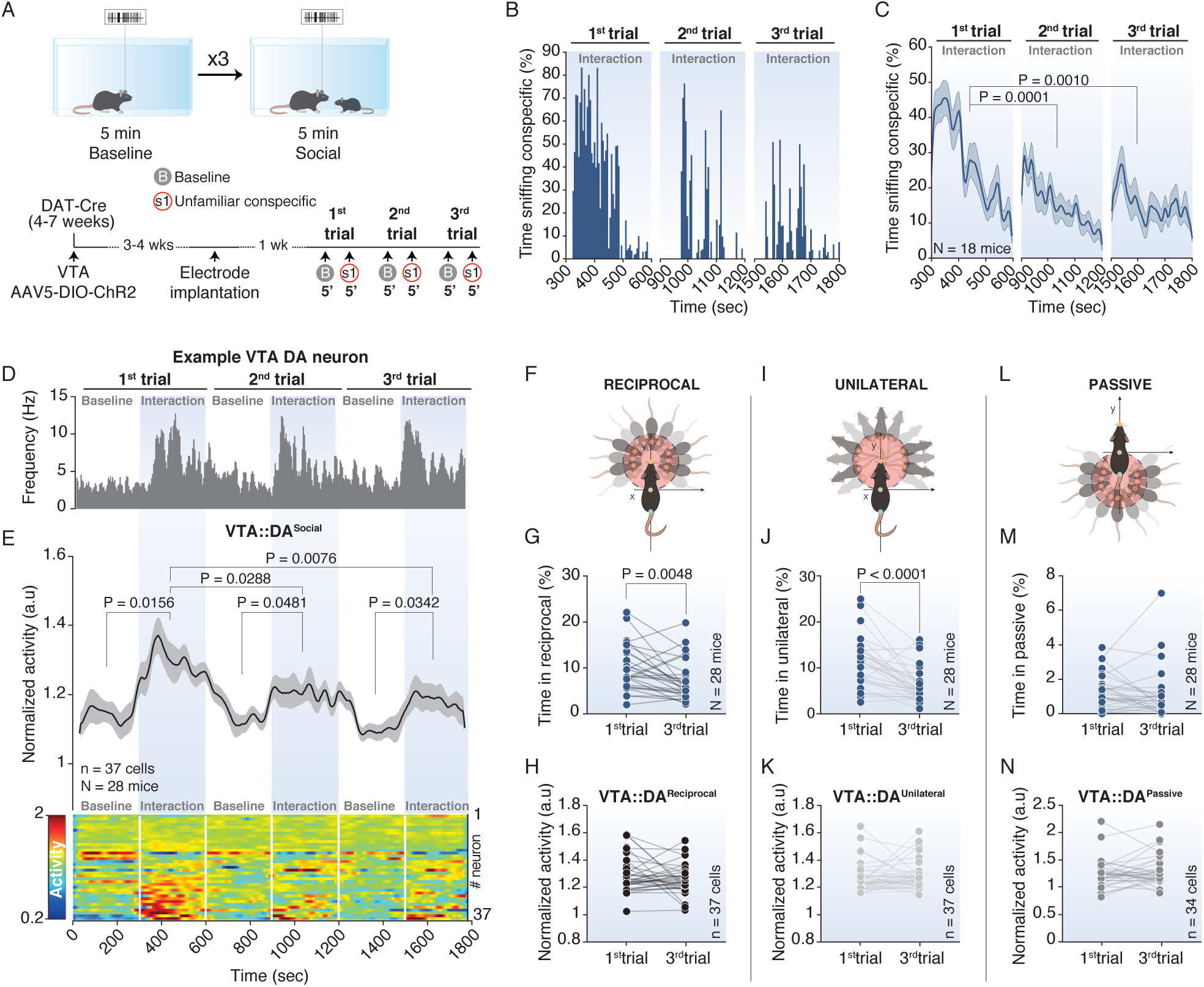
VTA DA neuron activity encodes active interaction value and adapts to social context. **(A)** Top: Schema of the repeated free social interaction task between. Down: Experimental time-course of the repeated free social interaction. **(B)** Example of time sniffing the social stimulus across the 3 trials for one experimental mouse. **(C)** Time sniffing the social stimulus during the 3 trials for all the experimental mice. Friedman test (χ^2^_(18)_ = 18.78, P < 0.0001) followed by Bonferroni-Holm correction. **(D)** Example of a VTA DA neuron frequency during repeated free social interaction across the 3 trials. **(E)** Top: Time course of the normalized VTA DA neuron activity during baseline and social session for the 3 trials. RM one-way ANOVA (Time main effect: F_(5,36)_ = 9.14, P < 0.0001) followed by Bonferroni-Holm correction. Down: Time course of the heatmap for the normalized VTA DA activity for each neuron across the 3 trials. **(F)** Schema of the reciprocal interaction. **(G)** Comparison of time in reciprocal interaction between 1^st^ and 3^rd^ trials. Wilcoxon test (W = -242). **(H)** Comparison of the normalized VTA DA activity during reciprocal interaction between 1^st^ and 3^rd^ trials. Wilcoxon test (W = -167, P = 0.2132). **(I)** Schema of the unilateral interaction. **(L)** Comparison of time in unilateral interaction between 1^st^ and 3^rd^ trials. Wilcoxon test (W = -346). **(M)** Comparison of the normalized VTA DA activity during unilateral interaction between 1^st^ and 3^rd^ trials. Wilcoxon test (W = 135, P = 0.3160Paired t-test (t_(36)_ = -2.126). **(N)** Schema of the passive interaction. **(O)** Comparison of time in passive interaction between 1^st^ and 3^rd^ trials. Wilcoxon test (W = -104, P = 0.2427). **(P)** Comparison of the normalized VTA DA activity during passive interaction between 1^st^ and 3^rd^ trials. Wilcoxon test (W = 2, P = 0.9926).

Within the VTA, a subset of DA neurons encodes the novelty of the rewards more than its value (*15*). Moreover, we have previously shown a progressive decrease in interaction during exposure time and following repeated exposure to the social stimulus (*16*). Therefore, we asked whether VTA DA neuron can differentially encode the value and the novelty associated with social interaction. In the free interaction task, we repeatedly exposed the experimental mouse to the same social stimulus (Fig. 2A). As previously shown, (*16*) time sniffing the social stimulus habituated within and between the 3 trials (Fig. 2B, C). Moreover, we observed habituation of the overall VTA DA activity (Fig. 2D, E) but not in the bursting nor the tonic activity between trials (Suppl. Fig. 4A, B) suggesting an adaptation of VTA DA neurons to the social context.

However, when we time locked the activity to the interaction bouts, we observed that, although the time spent in active contacts habituated, at population level, VTA DA activity did not adapt between sessions (Fig. 2F-N), showing an increased activity during active and passive contacts in the 3^rd^ trial (Suppl. Fig. 4C-Z). Remarkably, we observed high response heterogeneity between neurons throughout the sessions and contact types (Suppl. Fig. 5A-H). Specifically, while a smaller percentage of the neurons recorded responded to reciprocal interaction during the 3^rd^ trial, a higher percentage were recruited by unilateral interactions suggesting that neuronal activity encodes differentially the two active contacts. Thereby, based on our results, the data indicate that different DA neurons of the VTA are recruited to encode the novelty and the value of different social contacts.

The increase in VTA DA neuron activity when the mice engage in an active contact suggests a role of DA in motivation and learning within a social context. To investigate the reinforcing properties of social interaction (*17, 18*), we designed an instrumental task (Social instrumental task: SIT) using a two-chambered shuttle boxes divided by a gridded auto-guillotine door. The lever in one chamber controlled the opening of the door allowing a 7 sec interaction between two conspecificfics (Fig. 3A). We trained the experimental mice to associate the lever press with the door opening to gain access to a sex-matched social stimulus in a daily 20 minutes sessions. After 10 days of training (shaping phase), experimental mice were tested for 15 days (instrumental phase; Fig. 3B, C). The increased number of lever-presses (Fig. 3D-F) and transitions from the lever to the interaction zone (Fig. 3G, H), between the shaping and the instrumental phases, indicates that the majority of the mice learned the task (Suppl. Fig. 6A). During the instrumental phase, we observed an increase in the peak of velocity at the lever press (Fig. 3I-K). This occurred together with an increase in fast transitions (< 2sec) and a decrease of missed trials compared to the shaping phase, while slow and delayed transitions did not change (Fig. 3L). These results suggest that interaction with social stimulus induces reinforcement learning.

**Figure 3.**
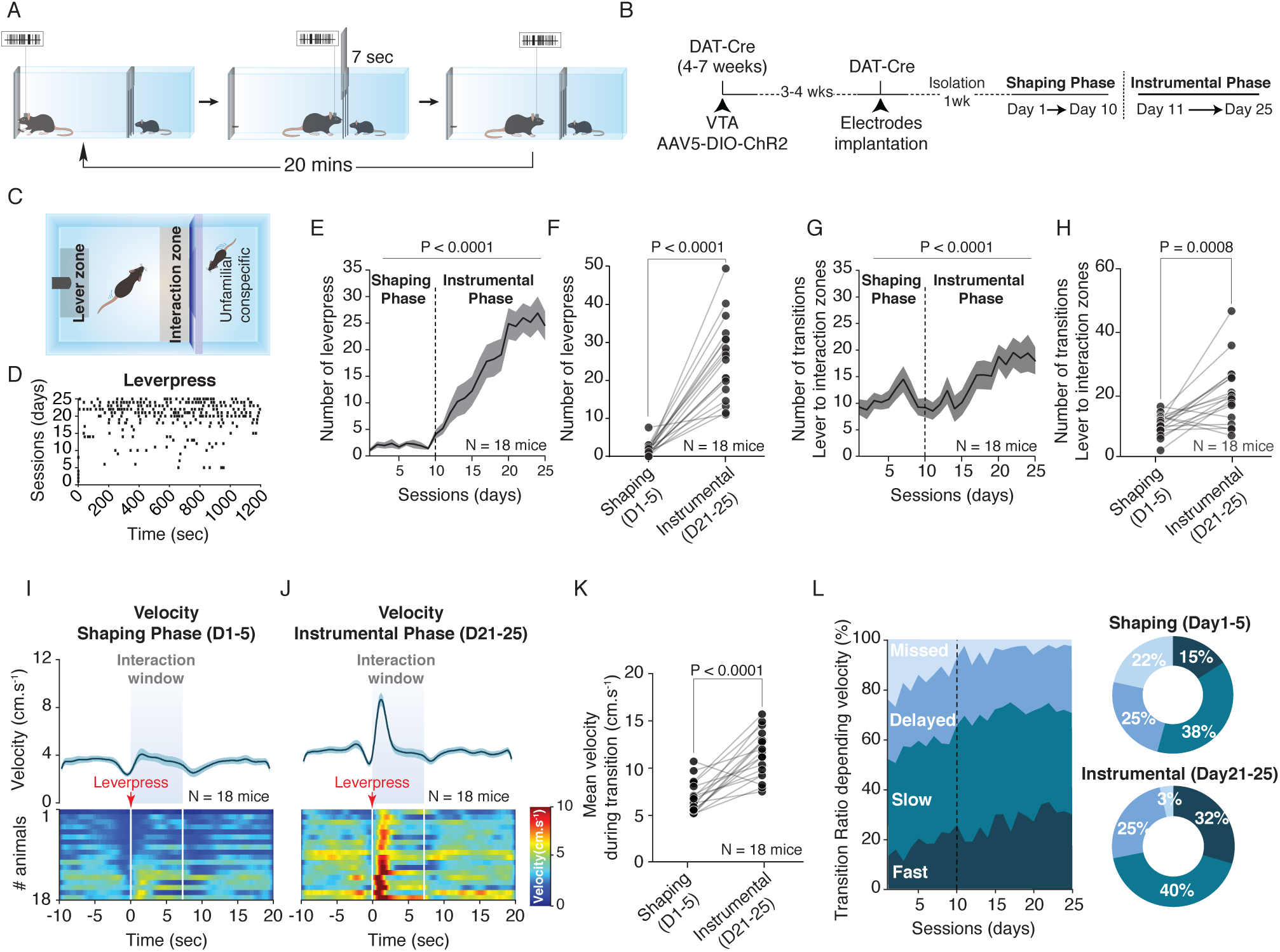
Social Instrumental Task (SIT). **(A)** Schema of the SIT for one session. **(B)** Experimental time-course of the SIT. The DAT-Cre mice are injected with an AAV-DIO-ChR2 and implanted with recordings electrodes in the VTA prior to perform the SIT. **(C)** Schema of the Operant Chamber defining the different zones (lever and interaction zones). **(D)** Raster plot example of lever-presses of 1 mouse across the different sessions in shaping and instrumental phases. **(E)** Number of lever-presses across different phases (shaping and instrumental). Repeated measure (RM) one-way ANOVA (Time main effect: F_(24,17)_ = 30.18, P < 0.0001). **(F)** Comparison of the number of lever-press between the shaping and the instrumental phases. Wilcoxon test (W = 0). **(G)** Number of transitions between the lever and the interaction zones. Friedman test (χ^2^_(18)_= 83.95, P < 0.0001). **(H)** Comparison of the number of transitions between the shaping and the instrumental phases. Paired t-test (t_(17)_ = 4.074). **(I)** Top: Peri-event time histogram (PETH) of the velocity for the shaping phase centered on the lever-presses. Down: PETH’s heatmap of the velocity for each animal during the shaping phase. **(J)** Top: PETH of the velocity for the instrumental phase centered on the lever-presses. Down: PETH’s heatmap of the velocity for each animal during the instrumental phase. **(K)** Comparison of the averaged velocity during the transition from the lever to the interaction zone, between the shaping and the instrumental phases. Paired t-test (t_(17)_ = 6.650). **(L)** Left: Ratio of the different transitions depending on the velocity across the sessions: fast, slow, delayed and missed. Right: Proportion of the different transitions between the shaping and the instrumental phases.

To test the goal directed properties of the task, we verified the experimental animal’s ability of extinguishing and reinstating operant responding in a different cohort of mice (Supp. Fig. 6B). Remarkably, the number of lever-presses (Suppl. Fig. 6C, D) and transitions (Suppl. Fig. 6E, F) decreased during the extinction, and increased back during the reinstatement phase. While the peak of velocity during the lever and interaction zones transitions increased in the instrumental and reinstatement phases, we did not notice a significant decrease during the extinction phase (Suppl. Fig. 6G, H). However, the proportion of fast transitions (< 2sec) between the lever and the interaction zones were increased during the instrumental and reinstatement phases compared to shaping and extinction phases (Suppl. Fig. 4I-J). Altogether, our data strongly suggest the reinforcing and motivational properties of social interaction with an unfamiliar conspecific.

We then recorded VTA DA neuron activity during the SIT. During the shaping phase, by averaging the trials across the recorded VTA DA neurons, at population level, we observed an overall increase in the spike frequency and bursting activity when the social stimulus was accessible, whether the neuronal activity was aligned on the lever-press (Fig. 4A-C, Suppl. Fig. 7B-E) or on the entry in the interaction zone (Suppl. Fig. 7A). Importantly, VTA DA neurons increased their activity at the onset of the lever-press (Fig. 4D-F), before the entry in the interaction zone (Suppl. Fig 7F), during the instrumental phase. On the contrary, burst activity remained elevated during the opening period of the door (Suppl. Fig. 7G-J).

**Figure 4.**
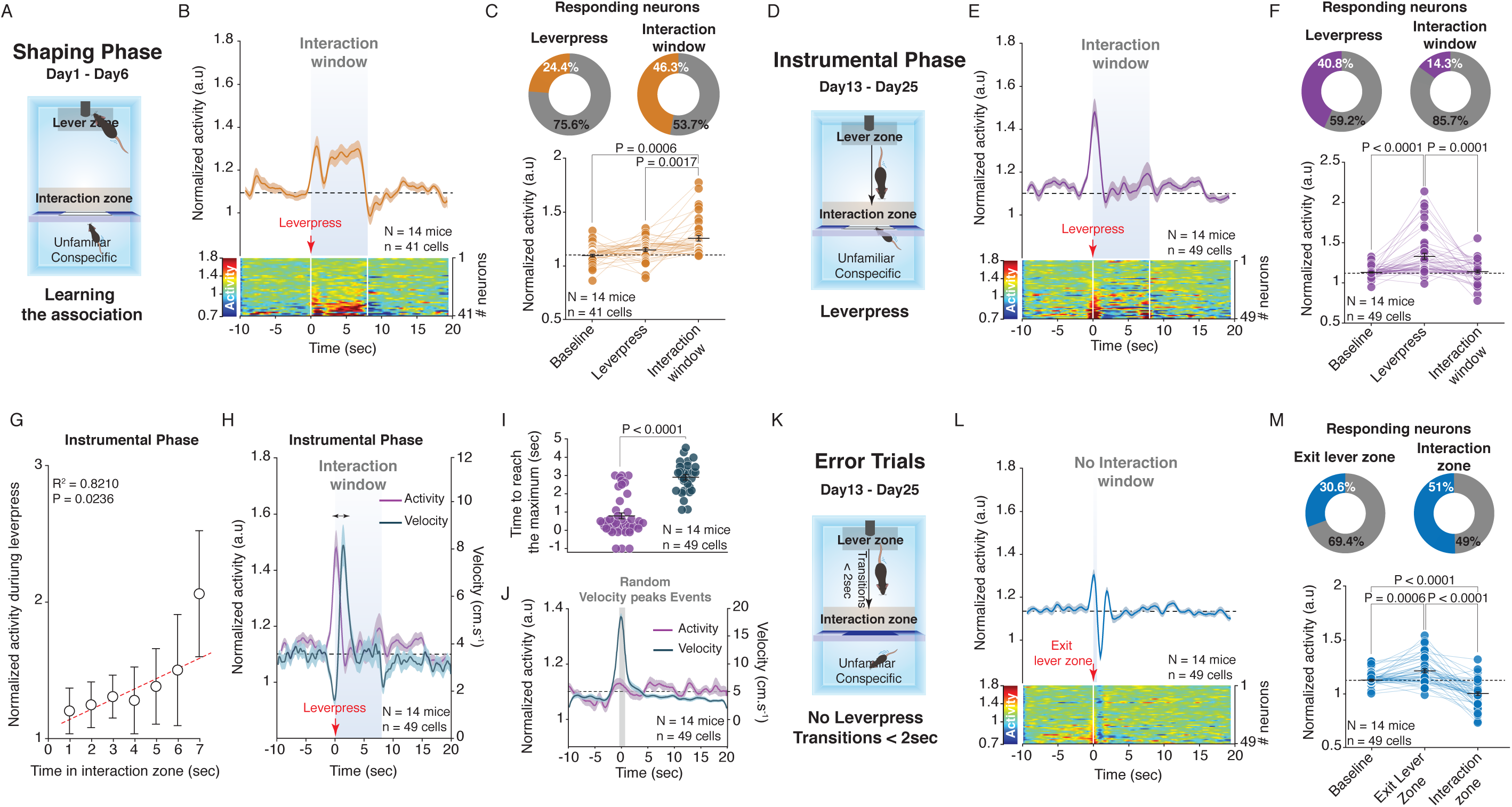
VTA DA neurons encode a social reward prediction error during the SIT. **(A)** Schema of the operant chamber defining the different zones (lever and interaction zones) during the shaping phase. **(B)** Top: PETH of the normalized VTA DA activity during the shaping phase centered on the lever-presses. Down: PETH’s heatmap of the normalized VTA DA activity for each neuron recorded. **(C)** Top: Proportion of the VTA DA neurons increasing their activity during the lever-press (Left) and the interaction window (Right). Down: Quantification of the normalized VTA DA activity during baseline (9 sec before the lever-press), lever-press (−1 and +1 sec around the lever-press) and the interaction window (6 last seconds of the interaction window). Friedman test (χ^2^_(41)_ = 21.66, P < 0.0001) followed by Bonferroni-Holm correction. **(D)** Schema of the operant chamber defining the different zones during the instrumental phase. **(E)** Top: PETH of the normalized VTA DA activity during the instrumental phase, centered on the lever-presses. Down: PETH’s heatmap of the normalized VTA DA activity for each neuron recorded. **(F)** Top: Proportion of the VTA DA neurons increasing their activity during the lever-press (Left) and the interaction window (Right). Down: Quantification of the normalized VTA DA activity during baseline, the lever-press and the interaction window. Friedman test (χ^2^_(49)_ = 32.37, P < 0.0001) followed by Bonferroni-Holm correction. **(G)** Normalized VTA DA activity during the lever-press in function of the consecutive time in the interaction zone after transition. Linear regression with Pearson’s correlation coefficient (R^2^ = 0.8210, P = 0.0236). **(H)** PETH of the normalized VTA DA activity (same data than in panel E) and velocity of the mice during the instrumental. The double arrow represents the delay between the VTA DA activity and the velocity peaks. **(I)** Quantification of the time to reach the maximum VTA DA activity or maximum velocity around the lever-press (1 sec before and 3 sec after). Mann-Whitney U test (U = 1814). **(J)** PETH centered to similar velocity peaks events higher than 20 cm.s^-1^ during the instrumental phase with the corresponding VTA DA activity. **(K)** Schema of the operant chamber with the different zones during error trials of the instrumental phase. **(L)** Top: PETH of the normalized VTA DA activity during the error trials, centered on the exit of the lever zone. Down: PETH’s heatmap of the normalized VTA DA activity for each neuron recorded. **(M)** Top: Proportion of the VTA DA neurons increasing their activity during the exit of the lever zone (Left) and decreasing activity following the transitions, in the interaction zone when the door is closed (Right). Down: Quantification of the normalized VTA DA activity during baseline, the exit of the lever zone and in the interaction zone. Friedman test (χ^2^_(49)_ = 61.27, P < 0.0001) followed by Bonferroni-Holm correction.

Further analysis showed correlation between the VTA DA activity at the moment of lever-press and the future time spent in the interaction zone (Fig. 4G). These results suggest that the VTA DA activity during the lever press predict the value of interaction. Remarkably, VTA DA activity was not directly correlated with locomotor activity. Indeed, the peak of VTA DA activity occurred before the peak of velocity, at the action initiation, but similar peaks of velocity, outside trials, during the instrumental phase are not preceded by a significant peak of neuronal activity (Fig. 3H-J). These results suggest that the activity of VTA DA neurons encode the positive value of social interaction through a phasic signal.

We observed transitions in which the experimental mice transited from the lever to the interaction zone in less than 2 seconds without pressing the lever. We considered these trials as error trials (Fig. 4K) and we recorded VTA DA neurons activity when the animals committed the error. Interestingly, the increased activity recorded when the mice were leaving the lever zone was followed by a decrease in VTA DA firing when the mice expected to interact with the social stimulus (Fig. 4L, M; Suppl. 8A-D). As control, we then compared the error trials with transitions made in less than 2 seconds occurring after a lever-press (correct trials) (Suppl. Fig. 8E). On these trials, when the activity was aligned on the exit from the lever zone, we still observed the increased activity of VTA DA neurons (Suppl. Fig. 8F, G). As for all the trials of the instrumental phase, we also observed an increase in bursting activity (Suppl. Fig. 8I-L). However, we revealed a significant difference between the error and correct trials (Suppl. Fig. 8H) by comparing the phasic increase at the exit of the lever zone and the decrease of VTA DA activity during the interaction window. In line with these observations, after instrumental phase, some mice underwent a random phase, consisting in trials with 50% probability to access the social stimuli (Suppl. Fig. 9A, B). When social stimulus was unexpectedly not accessible, a strong decrease of VTA DA activity was observed after the lever-press (Suppl. Fig. 9C, D). Altogether, these data indicate that VTA DA neurons encode Social Reward Prediction Error (SRPE), support social reinforcement learning and that social interaction has a motivational value *per se*.

Altogether our data indicate that VTA DA neuron activity represents the signal during the initial period of the active social contacts and, while individual neurons are recruited differentially, the interaction with conspecific does not adapt during repeated exposure. These findings suggest that neuronal processes underlying social interaction have specific characteristic that may be unique compared to other reward. Importantly, our findings support the role of DA in social reward learning through the Social Reward Prediction Error and strongly suggest that the initiation of the interaction toward the conspecific rather than the consumption, works as a learning signal and promotes motivational state in social behavior. Understanding of the neuronal mechanisms by which social reward modulates motivation will finally help to more clearly understand how deficits in prediction error in social context may lead to social deficits related to psychiatric disorders such as autism (*19*).

## Acknowledgments

We would like to thank Christian Luscher, Manuel Mameli, Philippe Faure, Jérémie Naudé and Sebastiano Bariselli for the comments to the manuscript. We would like also to thank Sebastien Pellat for the technical support.

## Author contributions

C.P.S., B.G. and C.B. conceived the project and wrote the manuscript. C.P.S., B.G. performed the electrophysiological recordings and the behavioural experiments with the help of B.R and M.T. C.P.S., B.G. performed all the analysis and statistics.

## Methods

### Animals

The study was conducted using wild type (WT) and transgenic mice with C57BL/6J background. WT mice were obtained from Charles River. For DA neuron-specific manipulations DAT-iresCre (Slc6a3^tm1.1(cre)Bkmn^/J, called DAT-Cre in the rest of manuscript) were employed. Only males’ animals were used for all the experiments conducted. Mice were housed in groups (weaning at P21 – P23) and isolated prior the experiments, under a 12 hours light – dark cycle (7:00 a.m.–7:00 p.m.). All physiology and behavior experiments were performed during the light cycle. For WT and DAT-Cre mice, multiple behavioral tests were performed with the same group of animals. All the procedures performed at UNIGE complied with the Swiss National Institutional Guidelines on Animal Experimentation and were approved by the respective Swiss Cantonal Veterinary Office Committees for Animal Experimentation.

### Multi-unit recording system – Microdrive

The VTA DA neurons recording was realized thanks to 2 octrodes, each constituted of 8 Nickel-Chrome (NiCr) coated wires of 15μm diameter, and a reference electrode for each octrode, constituted of stainless steel wires (110 μm). The octrodes are inserted in a homemade microdrive composed of a central piece containing a cannula as guide and a connector (Electrode interface board EIB, Neuralynx) where the recording and amplifier cable are plugged. After implantation, depth modulation is controlled by moving the central piece with a micro-screw. The 2 octrodes and an optic fiber are glued together through the cannula and implanted at the same time with a difference of 200 – 500μm between the tips of the octrodes and the optic fiber. Once the ensemble mounted, the impedance is uniformed (≈ 300 kOhms) at the octrodes tips with a diluted gold plating in a polyethylene glycol (C_2_H_4_O) solution.

The neuronal activity was recorded using Digital Lynx 4SX acquisition system, with 32kHz sampling rate (Neuralynx). A high band-pass filter (600Hz – 6000Hz) was applied during the recording to extract the fast electrical impulsions (the spikes).

### Surgery

Injection of rAAV5-Ef1*α*-DIO-hChR2(H134R)-eYFP (Titer ≥ 4.2×10^12^ vg.mL-1, UNC Vector Core) was performed in DAT-Cre mice at 4 – 7 weeks. Mice were anesthetized with a mixture of oxygen (1 L/min) and isoflurane 3% (Baxter AG, Vienna, Austria) and placed in a stereotactic frame (Angle One; Leica, Germany). The skin was shaved, locally anesthetized with 40 – 50 µL lidocaine 0.5% and disinfected. Unilateral craniotomy (1 mm in diameter) was then performed over the VTA at following stereotactic coordinates: ML ± 0.5 mm, AP −3.2 mm, DV −4.20 ± 0.05 mm from Bregma. The virus was injected via a glass micropipette (Drummond Scientific Company, Broomall, PA) into the VTA at the rate of 100 nl/min for a total volume of 500 nL. The implantation of the homemade Microdrive (see previous part) was then performed 2 weeks later using the same coordinates. Unilateral craniotomy was made above the VTA and bilateral craniotomy above the cerebellum to implant references wires. The Microdrive was then fixed on the skull using dental acrylic.

### Optogenetic photolabeling of VTA DA neurons

DAT-cre mice injected with rAAV5-Ef1*α*-DIO-hChR2(H134R)-eYFP and implanted with the microdrive underwent the optogenetic protocol to validate the dopaminergic nature of the neuron. When recorded neuron was suspected to be DA, based on the electrophysiological criteria (Firing < 12Hz, half-width spike > 1.5ms, regular spiking activity with typical bursting activity), mice were placed alone in a cage with bedding (20 × 30 × 15 cm). An optic fiber (homemade, materials from ThorLabs) was plugged and baseline neuronal activity was recorded during 90sec without stimulation. Then a 5Hz optical stimulation of blue laser (BioRay 488 nm 20 mW Elliptical Dot Laser, Coherent) protocol with a light-pulse duration of 5msec was applied with an expected power at the optic fiber tip of 8 – 12 mW (Master-8). After 1000 light-pulses the protocol was stopped, and the neuronal activity was still recorded during 1 min at baseline condition. Using the same procedure, a protocol of 20Hz light stimulation was then applied. The protocols were always applied after the different social experiments, at the end of the day, to avoid any influence of the light stimulation onto the tasks.

### Social Instrumental Task

The operant chamber (MedAssociates) is composed of 2 different compartments divided by a gridded auto-guillotine door: 1 chamber of 28 × 16 × 21 cm with the experimental DAT-Cre mouse and a lever-press on the wall opposing the door, and 1 chamber of 14 × 16 × 21 cm containing the social stimulus (unfamiliar sex-matched juvenile conspecific mouse C57BL/6J, 3–6 weeks). By pressing the lever, the gridded auto-guillotine door is opening during 7sec allowing the interaction between the experimental mouse and the social stimulus. The grid prevent the passage between chambers. After 7sec the door is closing. During the whole session, the experimental animals have the possibility to press without limit the lever to interact. The apparatus was cleaned using 70% ethanol after each session.

All the mice were isolated 1 week prior to the first session of the experiment to promote motivation to interact. The experimental mouse was placed in the corresponding chamber while neuronal recording was performed. To keep the same experimental conditions, the animals were always plugged even though neurons were not detected during the recording.

The task is performed with 1 session of 20 min per day and divided in several phases across the days:

- **Shaping phase (from Day 1 to Day 10):** The animals were trained to associate the lever-press with the opening of the door, and consequently, the social interaction. Every time the animals were in proximity of the lever, the experimenter, through the MedAssociates software, was opening the door. Every day the area around the lever was decreased and the last day of the shaping phase, the door was opened only when the mice were touching the lever.
- **Instrumental Phase (from Day 11 to Day 25):** The animals had to perform the task by themselves. Pressing only once on the lever was opening the door to interact with the conspecific.
- **Extinction Phase (from Day 26 to Day 75):** During this phase, the experimental mice were still able to press the lever and open the door, but no conspecific was present in the other chamber.
- **Reinstatement Phase (from Day 76 to Day 80):** An unfamiliar conspecific was reintroduced in the other chamber of the apparatus, and the experimental were able to interact with the stimulus by pressing the lever.
- **Random Phase (from Day 25 to Day 35):** After instrumental phase, if VTA DA neurons were still present, some animals underwent a paradigm where a lever-press was opening randomly the door with a 50% probability. Thereby the mice were not able to predict accurately the future social interaction with the unfamiliar conspecific.

The conspecific was changed every day to avoid experimental animals to interact twice with the same social stimulus during the different sessions of the task. During the instrumental phase, a mouse was considered as learner if it pressed at least 10 times the lever for 3 consecutives days. Otherwise the animal was considered as non-learner. The videos and neuronal recording were monitored and acquired using Neuralynx system. The animals were tracked and zones delimited using Ethovision software (Noldus).

At the end of the task, the animals were sacrificed. The viral infection and the recording electrode placement were verified.

### Free interaction task

DAT-Cre male mice, implanted with a Microdrive to record VTA DA neurons, performed a free interaction task. All the animals were isolated 1 week prior to the task. The mice were first placed in a cage like-homecage (20 × 40 × 10 cm) with bedding and VTA DA neuron activity was recorded. After 5 mins of neuronal activity recording, constituting the baseline activity, a social stimulus (unfamiliar conspecific sex-matched juvenile mouse) was introduced in the cage for 5 mins of free social interaction with the experimental mice. The baseline and social interaction conditions were repeated 3 times to obtain 3 different trials for a total duration of 30 mins. The same conspecific was used during the 3 different trials to study how the VTA DA neuron could adapt to a repeated social stimulus. At the end of each session of the task, the cage was cleaned using 70% ethanol.

The animals were tracked using DeepLabCut (*1*). Two models were built (experimental mice with implanted microdrive and stimuli mice) to detect nose, body and tail of each tracked subjects. Distance error between train and test dataset were of 0.0034cm for the two models. The rearing behavior was manually scored.

### Immunohistochemistry

DAT-Cre mice injected with rAAV5-Ef1*α*-DIO-hChR2(H134R)-eYFP were anesthetized with pentobarbital (Streuli Pharma) and sacrificed by intra-cardiac perfusion of 0.9% saline followed by 4% PFA (Biochemica). Brains were post-fixed overnight in 4% PFA at 4 °C. 24 hours later, they were washed with phosphate buffered saline (PBS) and then 50 μm thick sliced with a vibratome (Leica VT1200S).

Previously prepared slices were washed three times with PBS 0.1M. Brain slices were pre-incubated with PBS-BSA-TX buffer (10% BSA, 0.3% Triton X-100, 0.1% NaN_3_) for 60 minutes at room temperature in the dark. Subsequently, cells were incubated with primary antibodies diluted in PBS-BSA-TX (3% BSA, 0.3% Triton X-100, 0.1% NaN_3_) overnight at 4°C in the dark. The following day cells were washed three times with PBS 0.1M and incubated for 60 minutes at room temperature in the dark with the secondary antibodies diluted in PBS-Tween buffer (0.25% Tween-20). Finally, slices were mounted using Fluoroshield mounting medium with DAPI (abcam). In this study, the following primary antibody was used: rabbit polyclonal anti-Tyrosine Hydroxylase (1/500 dilution, abcam, ab6211). The following secondary antibody was used at 1/500 dilution: donkey anti-rabbit 555 (Alexa Fluor). Immunostained slices were imaged using the confocal laser scanning microscopes Zeiss LSM700 and LSM800. Larger scale images were taken with the widefield Axioscan.Z1 scanner.

### Analyses for In-vivo recording

#### Recording

All the *in-vivo* data acquired using Neuralynx system were extracted and analyzed offline using MatLab (The MathWorks). The spike-sorting was done using a custom MatLab code based on principal component analysis (PCA) and expectation-maximization of Gaussian mixture (EMGM). After spike-sorting procedure all the timestamps of the spikes as well as the voltage associated to each spike point were saved. Putative dopaminergic neurons (pDA) were first visually determined by wider waveform, slow firing pattern between 1 and 12Hz and typical triphasic bursting. Non-DAergic neurons (non-pDA) were determined by narrower waveform, high firing pattern or low firing pattern with burst event at high frequency (> 12Hz).

#### VTA neurons classification

Multiple electrophysiological features were extracted from the first 5min of each recording. For waveform features, spike total duration, half duration and half peak duration was used. For firing pattern features, burst and pause interspike interval (ISI) thresholds were determined and burst and pause strings were identified based on the Robust Gaussian Surprise (RGS) method (*2*). The probability distribution of each event (tonic, bursting, pause, spikes within tonic, spikes within burst, spikes within pause) were computed and the following properties of each distribution were extracted: mean, median, coefficient of variation, skewness and kurtosis. Finally, the frequency peak in the power spectrum was also extracted by using the Welch method to calculate power spectrum by averaging Fast Fourier Transforms of overlapping window divisions.

To check the possibility to classify VTA neurons based on electrophysiological properties, a cluster analysis was used based on the 58 features extracted. The features were first normalized by using zscore and by rescaling the values. Then, a PCA was processed and the number of components necessary to explain 99% of the variation was used to process a T-SNE. While the observation of two distinct cluster was evident, an EMGM was used to reliably quantify the membership of each recorded neurons in the two cluster. A threshold of 95% of probability of cluster membership was used to exclude neurons (1 pDA and 10 non-pDA). All the neurons identified by phototagging were present in the pDA cluster.

By using the MatLab classification learner app, different classification models were trained based on our dataset of 279 recorded neurons by using a 5 folds cross-validation. Ensemble of bagged classification trees reached the best performance with 99.6% of accuracy.

#### VTA DA activity and behavior analysis

For DA activity, the bursting activity was defined as described by Grace & Bunney (1984) (*3*): typically, a burst is starting when the inter-spike interval (ISI) is lower than 80msec, and is ending when the ISI is higher than 160msec. The activity outside the bursts were then considered as tonic activity.

The construction of Peri-event time histogram (PETH) was made by aligning and centering specific events. These events were obtained by coupling the Neuralynx digital acquisition system with others data acquisitions systems (such as Master-8 or MedAssociates operant chamber) that sent Time-to-Live (TTL) at specific times to link neuronal activity with events/stimuli or with events detected by synchronized video analysis of specific behaviors.

The neuronal recordings were binned depending the analysis time-window taken. For large time-scale (time-window > 600 sec), 1 sec bin was taken to average the spiking frequency. For low time-scale (time-window < 30 sec, such as PETH), 100 msec bin was taken to average the spiking frequency. Neuronal activity was normalized on the global averaged neuronal activity of a session. To get normalized spiking activity, the normalization was computed as following: 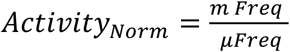; where *m Freq* is the averaged frequency of a given bin and *μ Freq* is the mean frequency of all the session. After normalization, a convolution using a Kernel-Gaussian sliding window of 16 bins was applied on the data (*gausswin* MatLab function).

The behavioral data associated with *in-vivo* recordings were 5sec binned and also smoothed using a Kernel-Gaussian sliding window of 3 bins.

#### Free Interaction

All the coordinates of the tracked position by DeepLabCut were corrected by using the body of the experimental mice as the origin coordinates and the nose of the experimental mice aligned to the y-axis. All the tracked position of the social stimulus was then plotted relatively to the position of the experimental mice. Coordinates with a likelihood lower than 95%, defined by DeepLabCut, were replaced by the last coordinates higher than 95% of likelihood. The distance and the angle between the experimental and the stimulus position were reported for each frame (40msec) and time distribution probability was computed. The neuronal activity was normalized as previously described when stimuli were present, then, for each frame, when the stimuli were closer than 20cm, the corresponding neuronal activity was reported and analyzed depending on the proximity and the angle in the same way than the time distribution of the stimuli position.

To extract interaction events, a proximity threshold of 5cm, an angle threshold from -110° to +110° (corresponding to the visual field of the experimental mice (*4*)) and a duration threshold from 0.2 to 2 seconds were applied regarding the position of the stimuli nose (active reciprocal interaction), the base of the stimuli tail (active unilateral interaction) or the base of the experimental mice tail (passive interaction). PETH analyzes were then performed and neuronal activity was aligned on the center of the defined interaction events and normalized as previously described using 10 seconds around each interaction event. To quantify the neuronal response, baseline activity was measured by the mean of the activity between 2 and 5 seconds before and after the event. The activity during the interaction was measured by the mean of activity 1sec before and 1sec after the interaction.

To determine responders and non-responders neurons, we calculated the p-value from t-tests for each neuron by comparing the baseline and interaction activity distributions from each trials. Every significant t-test determined if a neuron was responder or not. The average activity of the neuron during interaction (below or above the baseline) determined the positive or negative response. Neurons without response in any interaction were considered as non-responders and neurons with response in only a subset of interactions were considered as neutral for the interaction without response.

#### Social Instrumental Task

Data of position coordinates, velocity and transition events from lever to interaction zone of experimental mice were obtained directly from Ethovision software after the tracking. Lever-press timestamps event was obtained by TTL sent by MedAssociates apparatus. To analyze behavior and neuronal activity, further analyzes were performed to obtain different events. Transition performances during interaction window were defined as follow: fast (transitions < 2 sec), slow (2sec < transitions < 7 sec), delayed (7sec < transitions < 12 sec) and missed (transitions > 12 sec). Error trials events were defined by fast transitions not preceded by a lever-press and correct trials events by fast transitions preceded by a lever-press.

Velocity control events were defined by peak of velocity outside the trials (when door is closed) and higher than 20cm.s^-1^ (corresponding to the mean velocity of fast transitions when the door is open).

PETH analysis previously described was then performed. Velocity and/or neuronal activity were aligned:

- On lever-presses events, or entry in the interaction zone, during shaping phase between day 1 and day 6 and during instrumental phase between day 13 and day 25.
- For the false and correct trials, on the exit of the lever zone during fast transitions of the instrumental phase between day 13 and day 25. As the exit of the lever zone is not reaching the same temporal precision than a lever-press, the neuronal activity was then realigned to the lowest activity found 1 sec before or after the exit of the lever zone. For proper control, this readjustment was made in the same conditions for false and correct trials. VTA DA activity was then compared between both trials.
- For the random phase, on the lever-presses of the trials when the door was not opening.
- For control of the velocity, on the initiation of the action followed by a peak of velocity outside trials during instrumental phase between day 13 and day 25.

To analyze the reward prediction value, the mean of neuronal activity at the moment of the lever-press (between -1sec before and 1sec after) during the instrumental phase, between day 11 and day 20, was correlated with the time spent in the interaction zone.

By using the same approach than in the free interaction task, to determine responders and non-responders neurons, each trial of each neuron was compared between the baseline (10 seconds before the lever-press) and during the interaction window or lever-presses events (1 second before and after). We calculated the p-value from t-tests for each neuron by comparing the baseline and interaction window distributions. Every significant t-test determined if a neuron was responder or not.

### Statistical analysis

No statistical methods were used to predetermine the number of animals and cells, but suitable sample sizes were estimated based on previous experience and are similar to those generally employed in the field. The animals were randomly assigned to each group at the moment of viral infections or behavioral tests. Statistical analysis was conducted with MatLab (The Mathwork) and GraphPad Prism 7 (San Diego, CA, USA). Statistical outliers were identified by using the criterion *Mean*_*Value*_ ± 3 × *Std*_*Value*_ and excluded from the analysis. The normality of sample distributions was assessed with the Shapiro–Wilk criterion and when violated non-parametrical tests were used. When normally distributed, the data were analyzed with independent t test, paired t test, while for multiple comparisons one-way ANOVA and repeated measures (RM) ANOVA were used. When normality was violated, the data were analyzed with Mann–Whitney test, Wilcoxon matched-pairs signed rank test, while for multiple comparisons, Kruskal–Wallis or Friedman test were followed by Dunn’s test. For the analysis of variance with two factors (two-way ANOVA or RM two-way ANOVA), normality of sample distribution was assumed, and followed by Bonferroni-Holm correction test or Bonferroni post-hoc test. All the statistical tests adopted were two-sided. When comparing two samples distributions similarity of variances was assumed, therefore no corrections were adopted. Data are represented as the *Mean* ± *s. e. m*. and the significance was set at P < 0.05.

---

**Supplementary Figure 1.**
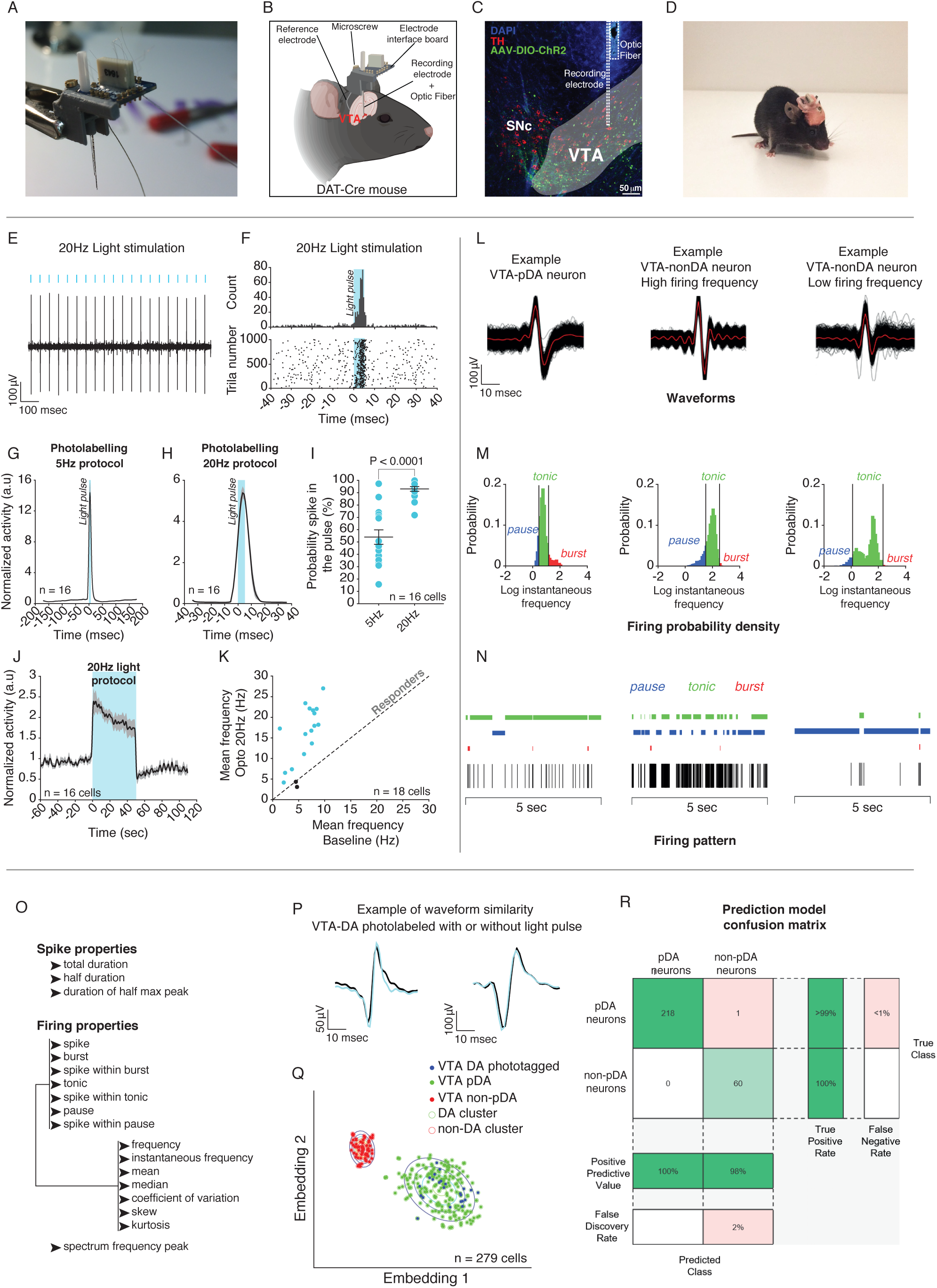
Recording, photolabeling and classification of the VTA DA neurons. **(A)** Picture of the microdrive implanted in DAT-Cre mice with recording electrodes and optic fiber. **(B)** Schema of implantation of the microdrive in the VTA. **(C)** Representative coronal image of immuno-staining experiments against Tyrosine Hydroxylase (TH) enzyme (in red) performed on midbrain slices of DAT-Cre adult mice infected with AAV5-Ef1α-DIO-ChR2-eYFP (green) in the VTA. **(D)** Picture of an implanted mouse for freely behaving recording. **(E)** Example trace of a VTA DA neuron following optogenetic stimulation at 20Hz. **(F)** Example PETH centered to the light pulse (top), of a VTA DA neuron responding to the 20Hz protocol stimulation with the corresponding raster plot for all the trials (down). **(G)** Averaged PETH, centered to the light pulse, for all the VTA DA neurons following the optogenetic protocol at 5Hz stimulation. **(H)** Averaged PETH, centered to the light pulse, for all the VTA DA neurons following the optogenetic protocol at 20Hz stimulation. **(I)** Probability to have a spike in the 10ms following the beginning of the light pulse for 5Hz and 20hz protocols. **(J)** Time course of all the photolabeled VTA DA neurons before, during and after the 20Hz protocol stimulation. **(K)** Mean frequency of the VTA DA neurons at baseline (without optogenetic stimulation) and during optogenetic stimulation at 20Hz. Responders are represented in blue while non-responders (example of 2 neurons) to the light stimulation are in black. **(L)** Waveforms examples of different neurons after spike-sorting for VTA-pDA (Left), VTA-nonDA high firing (Middle) and VTA-nonDA low-firing (Right) neurons. The red line represent the average of all the waveforms recorded during a session for one neuron. **(M)** Firing probability density for 3 different neurons (VTA-pDA, VTA-nonDA high firing and VTA-nonDA low-firing). This probability is the Log_10_ of the instantaneous frequency for a given neuron, and allows to extract general features such as tonic and bursting activity by using Robust Gaussian surprise method. **(N)** Examples of traces for the 3 different neurons with the tonic, bursting and pause activity. **(O)** Features used to separate all the neurons recorded independently of their nature. **(P)** Example of waveforms similarity between light stimulation and no-light stimulation of a photolabelled VTA DA neuron. **(Q)** Clustering classification of all the neurons recorded in the VTA. The neurons can be dopaminergic (VTA DA), putative dopaminergic (VTA pDA) or putative non dopaminergic (VTA non-pDA). Using tSNE analysis with all the features presented in panel (O), it is possible to separate two different cluters with the VTA non-pDA neurons on a side and the VTA DA and VTA pDA on the other side. All the phototagged neurons are included in the DA cluster. The VTA pDA (n = 1) outside the cluster and the VTA non-pDA (n = 10) included in the DAergic cluster are rejected from the analysis. **(R)** Confusion matrix of classification model (ensemble of bagged classification trees trained with 5 folds cross-validation; observations = 279; predictors = 58; accuracy = 99.6%).

**Supplementary Figure 2.**
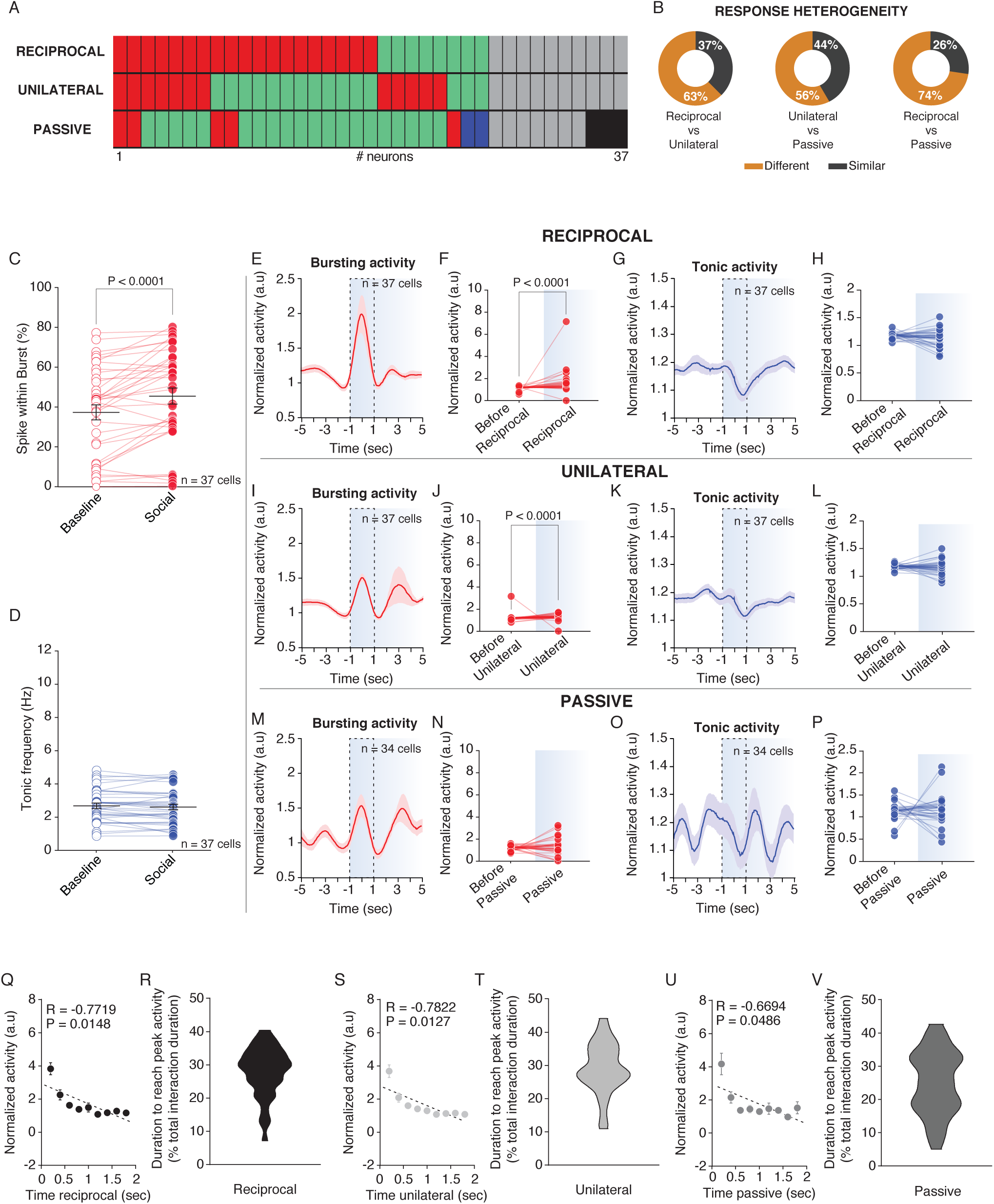
VTA DA bursting but not tonic activity increases during reciprocal and unilateral interactions. **(A)** Pattern of activity of the same VTA DA neurons depending on the type of interaction between reciprocal, unilateral and passive. **(B)** Response heterogeneity of VTA DA neurons between reciprocal, unilateral and passive interactions in the neurons responding to at least one type of interaction. **(C)** Spikes within burst of VTA DA neurons at baseline and during social interaction. Wilcoxon test (W = 96). **(D)** VTA DA tonic frequency at baseline and during social interaction. Paired t-test (t_(36)_ = 1.2307, P = 0.2264). **(E)** PETH of the normalized VTA DA bursting activity centered on the reciprocal interaction 5 seconds before and after the interaction. **(F)** Comparison between the VTA DA bursting activity before (5 seconds before) and during (−1 to 1 second around the interaction) active interaction. Wilcoxon test (W = 494). **(G)** PETH of the normalized VTA DA tonic activity centered on the active interaction 5 seconds before and after the interaction. **(H)** Comparison between the VTA DA tonic activity before (5 seconds before) and during (−1 to 1 second around the interaction) active interaction. Paired t-test (t_(36)_ = 1.702, P = 0.0974). **(I)** PETH of the normalized VTA DA bursting activity centered on the unilateral interaction 5 seconds before and after the interaction. **(J)** Comparison between the VTA DA bursting activity before (5 seconds before) and during (−1 to 1 second around the interaction) unilateral interaction. Wilcoxon test (W = 516, P < 0.0001). **(K)** PETH of the normalized VTA DA tonic activity centered on the unilateral interaction 5 seconds before and after the interaction. **(L)** Comparison between the VTA DA tonic activity before (5 seconds before) and during (−1 to 1 second around the interaction) unilateral interaction. Paired t-test (t_(36)_ = 0.8960, P = 0.3762). (M) PETH of the normalized VTA DA bursting activity centered on the passive interaction 5 seconds before and after the interaction. **(N)** Comparison between the VTA DA bursting activity before (5 seconds before) and during (−1 to 1 second around the interaction) passive interaction. Paired t-test (t_(36)_ = 0.7075, P = 0.4844). (O) PETH of the normalized VTA DA tonic activity centered on the passive interaction 5 seconds before and after the interaction. **(P)** Comparison between the VTA DA tonic activity before (5 seconds before) and during (−1 to 1 second around the interaction) passive interaction. Wilcoxon test (W = -29, P = 0.8132). **(Q)** Normalized VTA DA activity in function of the duration of bouts of reciprocal interaction. Pearson’s coefficient correlation. **(R)** Distribution of the duration to reach the peak of VTA DA activity during interactions (normalized in percentage of bouts duration). Mean = 28.00%, SEM = 1.242%. **(S)** Normalized VTA DA activity in function of the duration of bouts of unilateral interaction. Pearson’s coefficient correlation. **(T)** Distribution of the duration to reach the peak of VTA DA activity during interactions (normalized in percentage of bouts duration). Mean = 29.12%, SEM = 1.256%. **(U)** Normalized VTA DA activity in function of the duration of bouts of passive interaction. Pearson’s coefficient correlation. **(V)** Distribution of the duration to reach the peak of VTA DA activity during interactions (normalized in percentage of bouts duration). Mean = 24.96%, SEM = 1.596%.

**Supplementary Figure 3.**
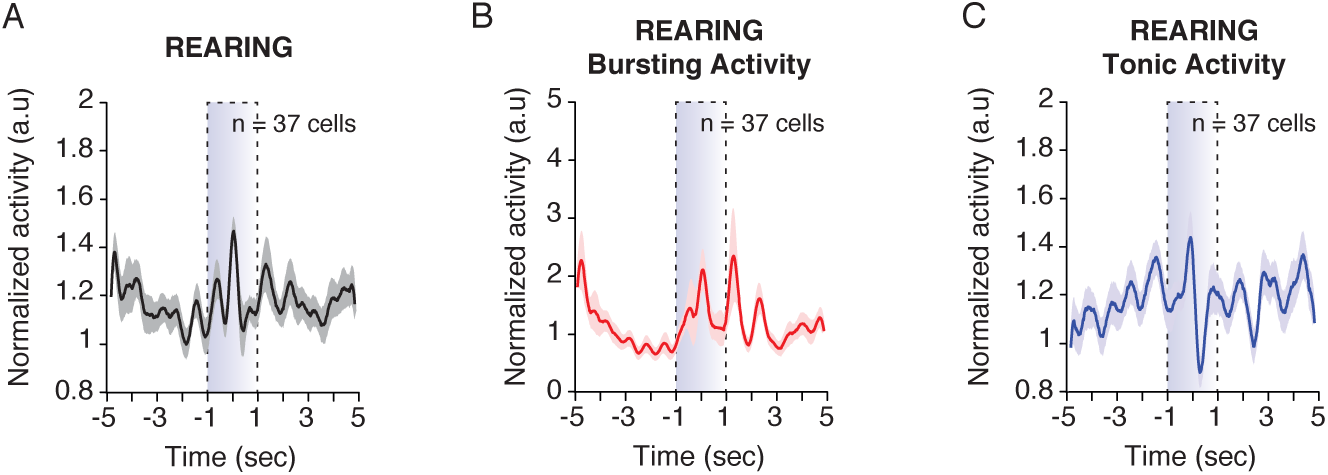
Rearing behavior does not induce VTA DA changes during free social interaction task. **(A)** PETH of the normalized VTA DA activity centered on the rearing behavior 5 seconds before and after the rearing. **(B)** PETH of the normalized VTA DA bursting activity centered on the rearing behavior 5 seconds before and after the rearing. **(C)** PETH of the normalized VTA DA tonic activity centered on the rearing behavior 5 seconds before and after the rearing.

**Supplementary Figure 4.**
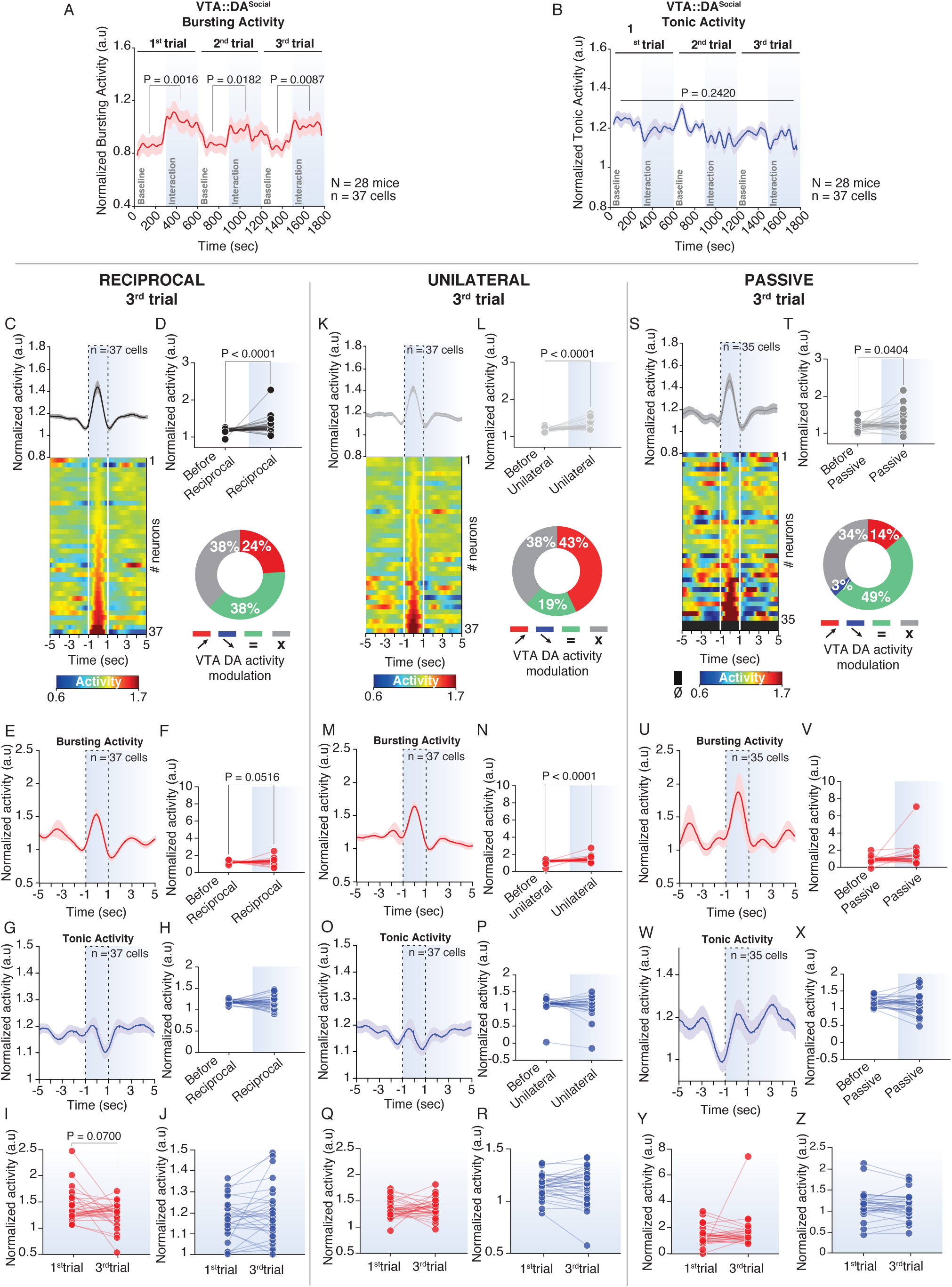
VTA DA activity during 3^rd^ trial of free interaction. VTA DA bursting and tonic activity do not adapt across repeated social exposure. **(A)** Time course of the normalized VTA DA bursting activity during baseline and social interaction for the 3 trials. RM one-way ANOVA (Time main effect: F_(5,36)_ = 12.42, P < 0.0001) followed by Bonferroni-Holm correction. **(B)** Time course of the normalized VTA DA tonic activity during baseline and social interaction for the 3 trials. RM one-way ANOVA (Time main effect: F_(5,36)_ = 1.20, P = 0.3095). (C) Normalized VTA DA activity during reciprocal interaction at 3^rd^ trial. Top: PETH of the normalized VTA DA activity centered on the reciprocal interaction 5 seconds before and after the interaction. Down: PETH’s heatmap of normalized VTA DA activity during reciprocal interaction for each neuron recorded. **(D)** Top: Comparison between the VTA DA activity before (5 seconds before) and during (−1 to 1 second around the interaction) reciprocal interaction. Wilcoxon test (W = 493, P < 0.0001). Down: Proportion of VTA DA neurons increasing (red) or not changing their activity (green) during reciprocal interaction. The grey labelling indicates non-responding neurons independently the type of interaction (reciprocal, unilateral or passive). **(E)** PETH of the normalized VTA DA bursting activity centered on the reciprocal interaction 5 seconds before and after the interaction during 3^rd^ trial. **(F)** Comparison between the VTA DA bursting activity before (5 seconds before) and during (−1 to 1 second around the interaction) reciprocal interaction. Wilcoxon test (W = 248, P = 0.0516). **(G)** PETH of the normalized VTA DA tonic activity centered on the reciprocal interaction 5 seconds before and after the interaction during 3^rd^ trial. **(H)** Comparison between the VTA DA tonic activity before (5 seconds before) and during (−1 to 1 second around the interaction) reciprocal interaction. Paired t-test (t_(36)_ = 1.088, P = 0.2839). (I) Comparison of the normalized VTA DA bursting activity during reciprocal interaction between 1^st^ and 3^rd^ trials. Wilcoxon test (W = -224, P = 0.0797). **(J)** Comparison of the normalized VTA DA tonic activity during reciprocal interaction between 1^st^ and 3^rd^ trials. Paired t-test (t_(36)_ = 1.706, P = 0.0966). **(K)** Normalized VTA DA activity during unilateral interaction at 3^rd^ trial. Top: PETH of the normalized VTA DA activity centered on the unilateral interaction 5 seconds before and after the interaction. Down: PETH’s heatmap of normalized VTA DA activity during unilateral interaction for each neuron recorded. **(L)** Top: Comparison between the VTA DA activity before (5 seconds before) and during (−1 to 1 second around the interaction) unilateral interaction. Wilcoxon test (W = 647). Down: Proportion of VTA DA neurons increasing, decreasing or not changing their activity during unilateral interaction. **(M)** PETH of the normalized VTA DA bursting activity centered on the unilateral interaction 5 seconds before and after the interaction during 3^rd^ trial. **(N)** Comparison between the VTA DA bursting activity before (5 seconds before) and during (−1 to 1 second around the interaction) unilateral interaction. Wilcoxon test (W = 566). **(O)** PETH of the normalized VTA DA tonic activity centered on the unilateral interaction 5 seconds before and after the interaction during 3^rd^ trial. **(P)** Comparison between the VTA DA tonic activity before (5 seconds before) and during (−1 to 1 second around the interaction) unilateral interaction. Wilcoxon test (W = -57, P = 0.6762). **(Q)** Comparison of the normalized VTA DA bursting activity during unilateral interaction between 1^st^ and 3^rd^ trials. (W = 122, P = 0.3461). **(R)** Comparison of the normalized VTA DA tonic activity during unilateral interaction between 1^st^ and 3^rd^ trials. Paired t-test (t_(36)_ = 0.3518, P = 0.7271). **(S)** Normalized VTA DA activity during passive interaction at 3^rd^ trial. Top: PETH of the normalized VTA DA activity centered on the passive interaction 5 seconds before and after the interaction. Down: PETH’s heatmap of normalized VTA DA activity during passive interaction for each neuron recorded. **(T)** Top: Comparison between the VTA DA activity before (5 seconds before) and during (−1 to 1 second around the interaction) passive interaction. Wilcoxon test (W = 250). Down: Proportion of VTA DA neurons increasing, decreasing or not changing their activity during passive interaction. **(U)** PETH of the normalized VTA DA bursting activity centered on the passive interaction 5 seconds before and after the interaction during 3^rd^ trial. **(V)** Comparison between the VTA DA bursting activity before (5 seconds before) and during (−1 to 1 second around the interaction) passive interaction. Wilcoxon test (W = 177, P = 0.1168). **(W)** PETH of the normalized VTA DA tonic activity centered on the passive interaction 5 seconds before and after the interaction during 3^rd^ trial. **(X)** Comparison between the VTA DA tonic activity before (5 seconds before) and during (−1 to 1 second around the interaction) passive interaction. Paired t-test (t_(34)_ = 0.8518, P = 0.4003). (Y) Comparison of the normalized VTA DA bursting activity during passive interaction between 1^st^ and 3^rd^ trials. (W = 23, P = 0.8236). **(Z)** Comparison of the normalized VTA DA tonic activity during passive interaction between 1^st^ and 3^rd^ trials. Wilcoxon test (W = -113, P = 0.3429).

**Supplementary Figure 5.**
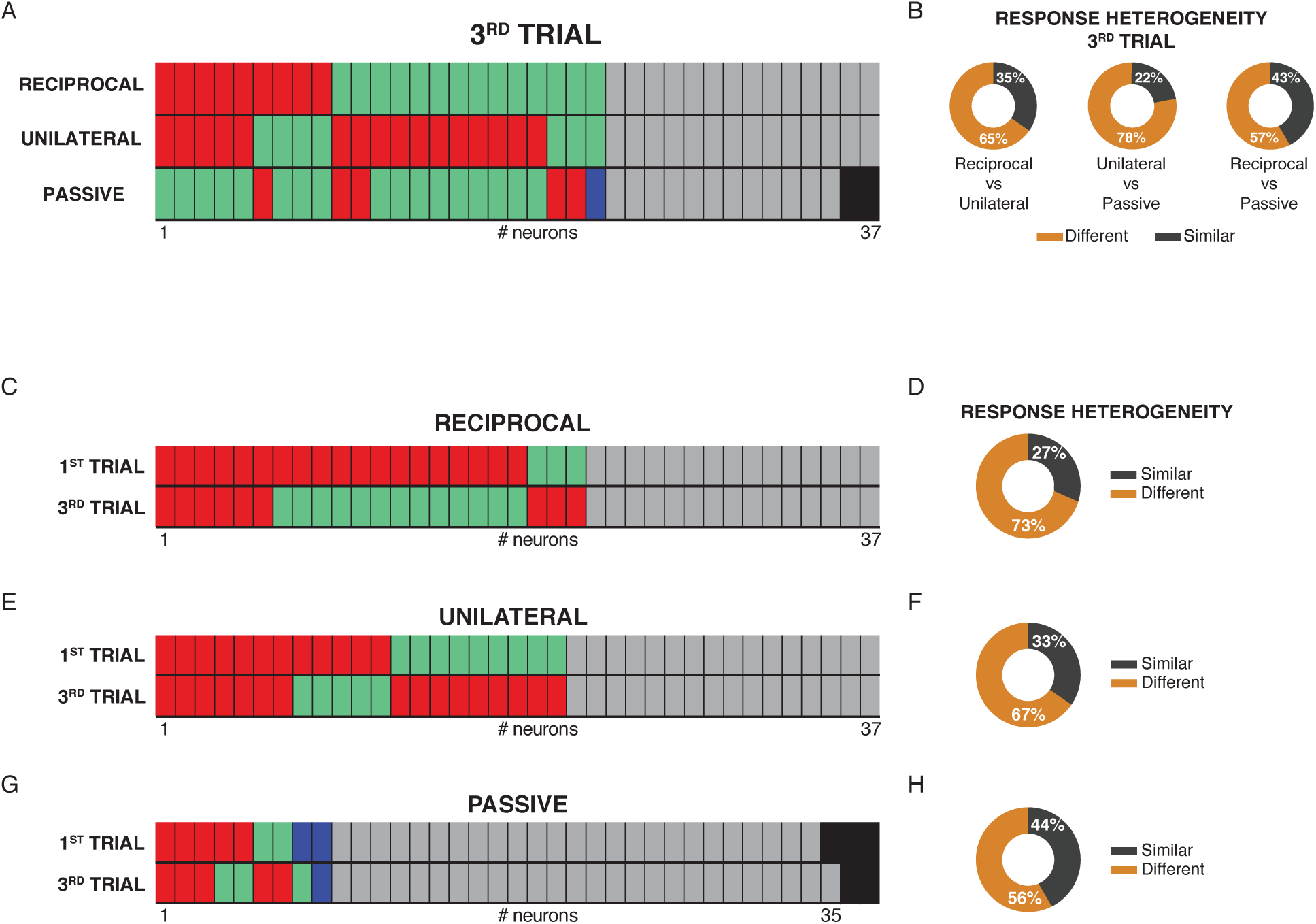
Heterogeneity of VTA DA activity depending on the type of contact and the exposure. **(A)** Pattern of activity of the same VTA DA neurons depending on the type of interaction between reciprocal, unilateral and passive during the 3^rd^ trial. **(B)** Response heterogeneity of VTA DA neurons between reciprocal, unilateral and passive interactions in the neurons responding to at least one type of interaction during the 3^rd^ trial. **(C)** Pattern of activity of the same VTA DA neurons depending on the trials of interaction for reciprocal contacts. **(D)** Response heterogeneity of the VTA DA neurons during reciprocal interactions in the neurons responding to at least one trial between the 1^st^ and 3^rd^ trials. **(E)** Pattern of activity of the same VTA DA neurons depending on the trials of interaction for unilateral contacts. **(F)** Response heterogeneity of the VTA DA neurons during unilateral interactions in the neurons responding to at least one trial between the 1^st^ and 3^rd^ trials. **(G)** Pattern of activity of the same VTA DA neurons depending on the trials of interaction for passive contacts. **(H)** Response heterogeneity of the VTA DA neurons during passive interactions in the neurons responding to at least one trial between the 1^st^ and 3^rd^ trials.

**Supplementary Figure 6.**
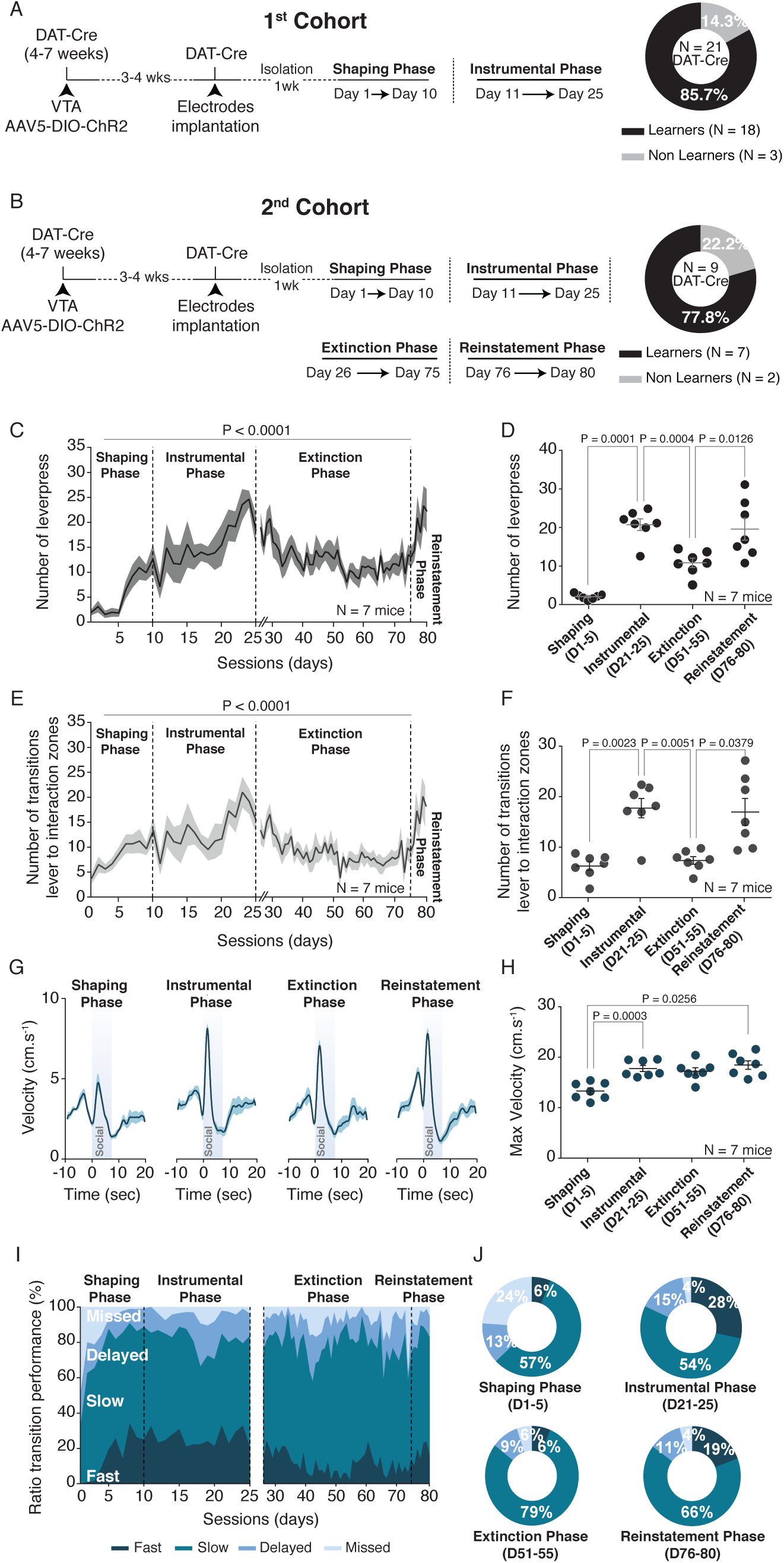
Extinction and reinstatement of the SIT. **(A)** Left: Experimental time-course of the Social Instrumental Task for the 1^st^ cohort (same cohort than in figure 1). The DAT-Cre mice are first injected with an AAV5-DIO-ChR2 and implanted with recordings electrodes in the VTA prior to perform the shaping and instrumental phases. Right: Proportion of learners and non-learners in the task for the 1^st^ cohort. **(B)** Left: Experimental time-course of the Social Instrumental Task for the 2^nd^ cohort. The DAT-Cre mice are first injected with an AAV5-DIO-ChR2 and implanted with recordings electrodes in the VTA prior to perform the shaping, instrumental, extinction and reinstatement phases of the SIT. Right: Proportion of learners and non-learners in the task for the 2^nd^ cohort. **(C)** Number of lever-presses across the days and the 4 different phases (shaping, instrumental, extinction and reinstatement) for the 2^nd^ cohort. RM one-way ANOVA (Time main effect: F_(79,6)_ = 4.72, P < 0.0001). **(D)** Comparison of the number of lever-press between the shaping (day1 – day5), instrumental (day21 – day25), extinction (day51 – day55) and reinstatement (day76 – day80) phases. RM one-way ANOVA (Phases main effect: F_(3,6)_ = 34.67, P < 0.0001) followed by Bonferroni-Holm correction. **(E)** Number of transitions between the lever and social zones for the 2^nd^ cohort. Friedman test (χ^2^ _(7)_= 176.5, P < 0.0001). **(F)** Comparison of the number of transitions between the shaping (day1 – day5), instrumental (day21 – day25), extinction (day51 – day55) and reinstatement (day76 – day80) phases. RM one-way ANOVA (Phases main effect: F_(3,6)_ = 18.69, P = 0.0001) followed by Bonferroni-Holm correction. (G) Peri-event time histogram (PETH) of the velocity for the shaping (day1 – day5), instrumental (day21 – day25), extinction (day51 – day55) and reinstatement (day76 – day80) phases, centered on the lever-presses 10 seconds before and 20 seconds after the lever-press. **(H)** Comparison of the maximum velocity during the transitions for all the different phases of the SIT. RM one-way ANOVA (Phases main effect: F_(3,6)_ = 6.557, P = 0.0106) followed by Bonferroni-Holm correction. **(I)** Ratio of the different transitions depending the velocity across the sessions of the different phases: fast (transitions < 2 sec), slow (2sec < transitions < 7 sec), delayed (7sec < transitions < 12 sec) and missed (transitions > 12 sec). **(I)** Proportion of the different transitions between shaping (day1 – day5), instrumental (day21 – day25), extinction (day51 – day55) and reinstatement (day76 – day80) phases.

**Supplementary Figure 7.**
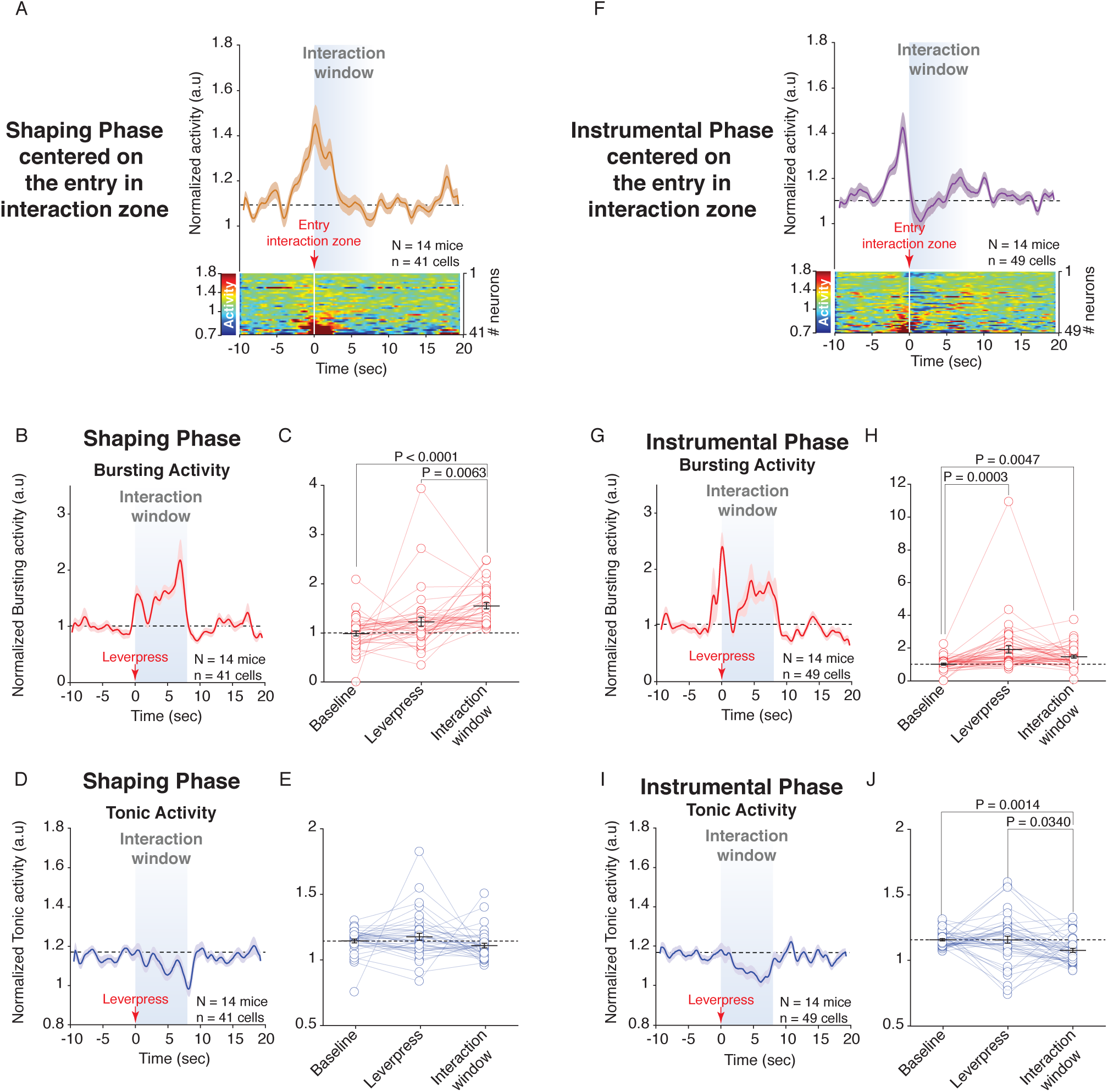
VTA DA bursting and tonic activity during shaping and instrumental phases. **(A)** Top: PETH of the normalized VTA DA activity during the shaping phase, centered on the entry in the interaction zone 10 seconds before and 20 seconds after the entry. Down: PETH’s heatmap of the normalized VTA DA activity for each neuron recorded during the shaping phase. **(B)** PETH of the normalized VTA DA bursting activity during the shaping phase, centered on the lever-presses 10 seconds before and 20 seconds after the lever-press. **(C)** Quantification of the normalized VTA DA bursting activity during baseline (9 seconds before the lever-press), lever-press and the interaction window. Friedman test (χ^2^_(41)_= 21.05, P < 0.0001) followed by Bonferroni-Holm correction. **(D)** PETH of the normalized VTA DA tonic activity during the shaping phase, centered on the lever-presses 10 seconds before and 20 seconds after the lever-press. **(E)** Quantification of the normalized VTA DA tonic activity during baseline (5 seconds before the lever-press), lever-press and the interaction window. Friedman test (χ^2^_(41)_= 7, P = 0.0302) followed by Bonferroni-Holm correction. **(F)** Top: PETH of the normalized VTA DA activity during the instrumental phase, centered on the entry in the interaction zone 10 seconds before and 20 seconds after the entry. Down: PETH’s heatmap of the normalized VTA DA activity for each neuron recorded during the instrumental phase. **(G)** PETH of the normalized VTA DA bursting activity during the instrumental phase, centered on the lever-presses 10 seconds before and 20 seconds after the lever-press. **(H)** Quantification of the normalized VTA DA bursting activity during baseline (9 seconds before the lever-press), lever-press and the interaction window. Friedman test (χ^2^_(49)_= 19.28, P < 0.0001) followed by Bonferroni-Holm correction. **(I)** PETH of the normalized VTA DA tonic activity during the instrumental phase, centered on the lever-presses 10 seconds before and 20 seconds after the lever-press. **(J)** Quantification of the normalized VTA DA tonic activity during baseline (9 seconds before the lever-press), lever-press and the interaction window. Friedman test (χ^2^_(49)_ = 15.49, P = 0.0004) followed by Bonferroni-Holm correction.

**Supplementary Figure 8.**
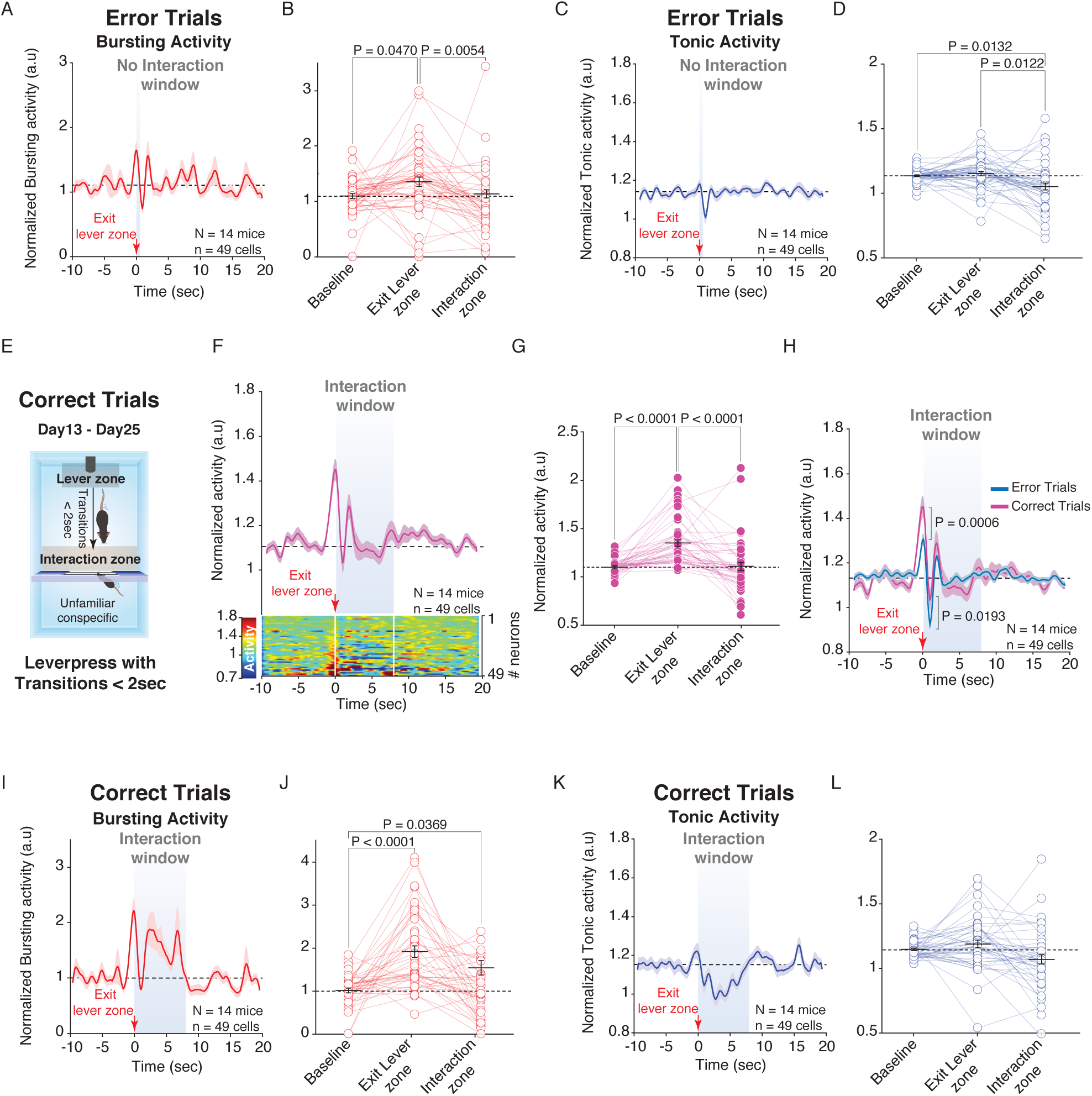
VTA DA bursting and tonic activity during error trials and comparison with correct trials. **(A)** PETH of the normalized VTA DA bursting activity during error trials, centered on the exit of the lever zone 10 seconds before and 20 seconds after the exit. **(B)** Quantification of the normalized VTA DA bursting activity during baseline (9 seconds before the exit), exit of the lever zone and in the interaction zone when the door is closed. Friedman test (χ^2^_(49)_ = 22.17, P < 0.0001) followed by Bonferroni-Holm correction. **(C)** PETH of the normalized VTA DA tonic activity during error trials, centered on the exit of the lever zone 10 seconds before and 20 seconds after the exit. **(D)** Quantification of the normalized VTA DA tonic activity during baseline (9 seconds before the exit), exit of the lever zone and in the interaction zone when the door is closed. Friedman test (χ^2^_(49)_ = 25.02, P < 0.0001) followed by Bonferroni-Holm correction. **(E)** Schema of the Operant Chamber defining the different zones (lever and interaction zones) during correct trials of the instrumental phase. **(F)** Top: PETH of the normalized VTA DA activity during the correct trials, centered on the exit of the lever zone 10 seconds before and 20 seconds after the exit. Down: PETH’s heatmap of the normalized VTA DA activity for each neuron recorded during the correct trials. **(G)** Quantification of the normalized VTA DA activity during baseline, the exit of the lever zone and in the interaction zone when the door is open, after the transition. Friedman test (χ^2^_(49)_= 44.80, P < 0.0001) followed by Bonferroni-Holm correction. **(H)** PETH of the normalized VTA DA activity during the correct and error trials (same data than figure 2, blue line), centered on the exit of the lever zone 10 seconds before and 20 seconds after the exit. Direct comparison of the VTA DA activity during the exit of the lever zone and in the interaction zone when the door is open (correct trials) or closed (error trials). Mann-Whitney U test (U = 1965). **(I)** PETH of the normalized VTA DA bursting activity during correct trials, centered on the exit of the lever zone 10 seconds before and 20 seconds after the exit. **(J)** Quantification of the normalized VTA DA bursting activity during baseline (9 seconds before the exit), exit of the lever zone and in the interaction zone when the door is open. Friedman test (χ^2^_(49)_= 28.35, P < 0.0001) followed by Bonferroni-Holm correction. **(K)** PETH of the normalized VTA DA tonic activity during correct trials, centered on the exit of the lever zone 10 seconds before and 20 seconds after the exit. **(L)** Quantification of the normalized VTA DA tonic activity during baseline (9 seconds before the exit), exit of the lever zone and in the interaction zone when the door is open. Friedman test (χ^2^_(49)_= 5.64, P = 0.0597).

**Supplementary Figure 9.**
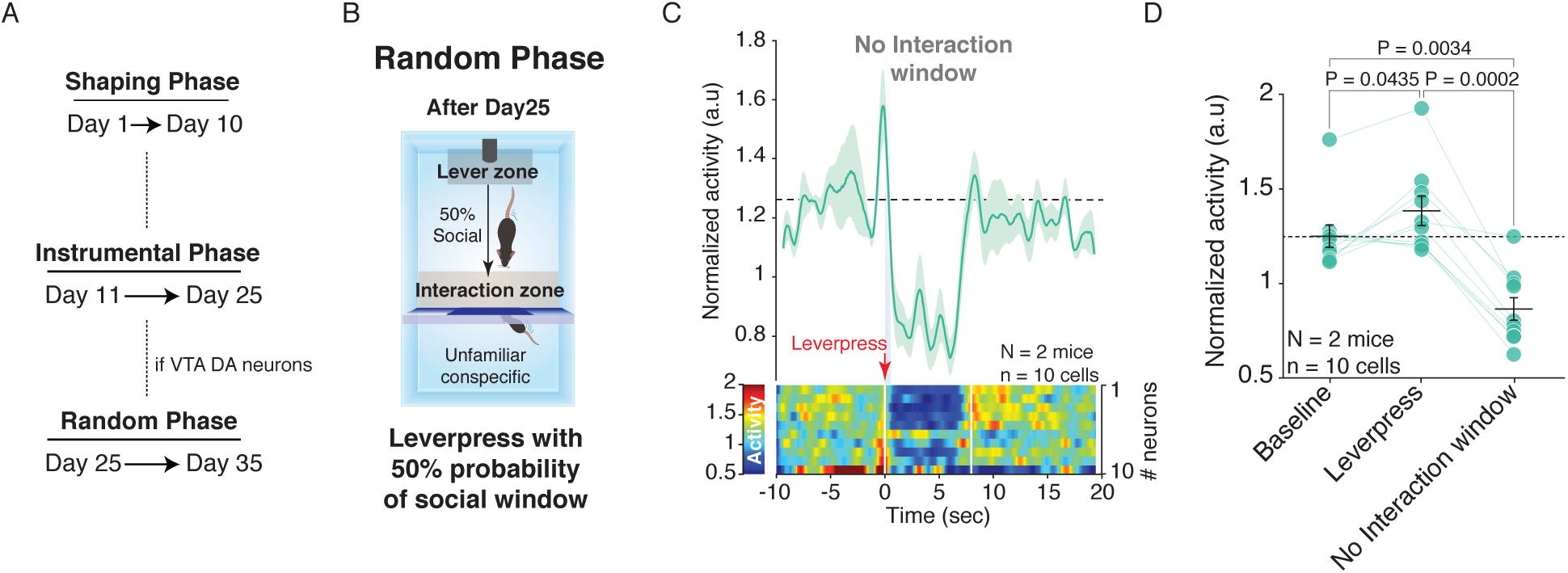
VTA DA activity during random phase. **(A)** Time course of the experimental design with the shaping, instrumental and random phases. **(B)** Schema of the Operant Chamber defining the different zones (lever and interaction zones) during the random phase. **(C)** Top: PETH of the normalized VTA DA activity during the random phase, centered on the lever-presses 10 seconds before and 20 seconds after the lever-press when the door stays closed. Down: PETH’s heatmap of the normalized VTA DA activity for each neuron recorded during the random phase. **(D)** Quantification of the normalized VTA DA activity during baseline (9 seconds before the lever-press), lever-press (1 second before and 1 second after the lever-press) and when the interaction window should has been occurred (6 last seconds of the expected social window). Friedman test (χ^2^_(10)_ = 12.6, P = 0.0018) followed by Bonferroni-Holm correction.

